# Propagating patterns of intrinsic activity along macroscale gradients coordinate functional connections across the whole brain

**DOI:** 10.1101/2020.12.11.422071

**Authors:** Behnaz Yousefi, Shella Keilholz

## Abstract

The intrinsic activity of the human brain, observed with resting-state fMRI (rsfMRI) and functional connectivity, exhibits macroscale spatial organization such as resting-state networks (RSNs) and functional connectivity gradients (FCGs). Dynamic analysis techniques have shown that the time-averaged maps captured by functional connectivity are mere summaries of time-varying patterns with distinct spatial and temporal characteristics. A better understanding of these patterns might provide insight into aspects of the brain’s intrinsic activity that cannot be inferred by functional connectivity, RSNs or FCGs. Here, we describe three spatiotemporal patterns of coordinated activity across the whole brain obtained by averaging similar ~20-second-long segments of rsfMRI timeseries. In each of these patterns, activity propagates along a particular macroscale FCG, simultaneously across the cortical sheet and in most other brain regions. In some areas, like the thalamus, the propagation suggests previously-undescribed FCGs. The coordinated activity across areas is consistent with known tract-based connections, and nuanced differences in the timing of peak activity between brain regions point to plausible driving mechanisms. The magnitude of correlation within and particularly between RSNs is remarkably diminished when these patterns are regressed from the rsfMRI timeseries, a quantitative demonstration of their significant role in functional connectivity. Taken together, our results suggest that a few recurring patterns of propagating intrinsic activity along macroscale gradients give rise to and coordinate functional connections across the whole brain.

## Introduction

The intrinsic activity of the brain is widely studied with resting-state functional magnetic resonance imaging (rsfMRI) and functional connectivity (Biswal et al., 1995). Commonly calculated as the Pearson correlation between rsfMRI timeseries from different pairs of brain areas, functional connectivity reflects aspects of the brain’s spatial organization. For instance, based on the functional connectivity profile of each area and similarity of profiles across areas, the brain can be parcellated into macroscale resting-state networks (RSNs) (Buckner et al., 2011; Yeo et al., 2011; Buckner and DiNicola, 2019) or described by a few macroscale functional connectivity gradients (FCGs) (Margulies et al., 2016; Marquand et al., 2017; Guell et al., 2018; Vos De Wael et al., 2018). These RSNs and FCGs have links to task-related activity and behavioral measures (Fox and Raichle, 2007; Smith et al., 2009; Smith et al., 2015; Margulies et al., 2016; Marquand et al., 2017; Guell et al., 2018), evidence not only for their neuronal basis but also that the spatial organization of the intrinsic activity serves a fundamental purpose.

Despite the success and prevalence of functional connectivity-based analyses, the resulting time-averaged spatial maps provide a limited window into the rich dynamics of the brain’s intrinsic activity. More recent analysis techniques that are sensitive to time-varying activity (e.g., windowed analysis (Keilholz et al., 2013; Allen et al., 2014), coactivation patterns (Petridou et al., 2013; Liu et al., 2018a), Hidden Markov Models (Chen et al., 2016; Vidaurre et al., 2017)) have convincingly argued, and in some cases, quantitatively demonstrated that functional connectivity is a mere summary of recurring patterns of activity with distinct spatial and temporal characteristics. A better understanding of the spatiotemporal characteristics of these patterns might provide insight into aspects of the brain’s intrinsic activity that cannot be gained by functional connectivity, RSNs or FCGs. The consideration of time-varying features of intrinsic activity explains behavioral variability in task performance and improves sensitivity of biomarkers to clinical alterations (Hutchison et al., 2013; Calhoun et al., 2014; Keilholz et al., 2017; Preti et al., 2017; Thompson, 2018), an indication that the time-varying features can capture additional important aspects of the underlying intrinsic brain dynamics.

One of the recurring spatiotemporal patterns of activity is the quasiperiodic pattern (QPP), obtained by identifying and averaging similar segments of rsfMRI timeseries (Majeed et al., 2011, 2009). The primary QPP (QPP1) in humans is approximately 20 seconds in duration and involves a cycle of activation and deactivation of different brain areas with various phases; most prominently, the cortical areas recognized as the task positive network (TPN) exhibit the opposite phase relative to the cortical areas known as the task negative or default mode network (DMN) (Majeed et al., 2011; Yousefi et al., 2018; Abbas et al., 2019a, 2019b; Briend et al., 2020). Focal propagation of activity is also observed, for example, along the medial prefrontal cortex (Majeed et al., 2011). The stereotyped activity represented by QPP1 recur frequently during rsfMRI and is remarkably similar across individuals (Yousefi et al., 2018). As expected, given the documented anticorrelation between the TPN and DMN in functional connectivity-based studies (Fox and Raichle 2007), QPP1 contributes substantially to functional connectivity within and between these networks (Abbas et al., 2019a, 2019b).

To better understand the spatiotemporal characteristics of QPP1 as a contributor to functional connectivity, we built upon our recent method improvements (Yousefi et al., 2018) to obtain the group QPP1 across the whole brain, utilizing the high quality and large dataset of the Human Connectome Project (HCP) (Van Essen et al., 2013). *First*, we inspected whether the propagation in QPP1 across the cerebral cortex has a principled relationship with the primary FCG (FCG1). FCG1 involves gradual changes in the similarity of functional connectivity profiles and is related to geodesic distance between areas across the cortical sheet. The TPN and DMN are situated with opposite signs along FCG1 (Margulies et al., 2016), consistent with their anticorrelation during QPP1. *Second*, we examined whether QPP1 reveals time-locked activity in the non-cortical regions of the brain, which could also propagate consistent with the reported FCGs in the cerebellum (Guell et al., 2018), hippocampus (Vos De Wael et al., 2018) and striatum (Marquand et al., 2017). *Third*, we explored the differences in the time of peak activity between the brain regions during QPP1 for suggestions about the driving mechanisms. *Fourth*, we developed a method to detect additional QPPs, and examined them for principled propagation across the brain consistent with other FCGs. *Finally*, we quantified the extent of the contribution of the first few QPPs to functional connectivity, the basis for RSNs and FCGs. Our results provide novel insights into the brain’s intrinsic activity, revealing aspects of whole-brain coordination that are only partially captured by RSNs and FCGs.

## Methods

We built upon our recent method improvements (Yousefi et al., 2018) and developed new approaches for this work, described broadly in the text that follows. All required details can be found in the supplementary materials (diagrams, table and additional text). Analyses of robustness to various methodological choices throughout are included at the end.

### Data and further preprocessing

We used the minimally preprocessed gray ordinate and FIX de-noised rsfMRI scans of the HCP S900 dataset (Glasser et al., 2013) and included all 817 individuals with four complete scans (repetition time (TR) of 0.72s, ~15 minutes per scan). The following preprocessing steps were additionally applied to each scan (Fig.S1). The grayordinate timeseries (~62K cortical vertices and ~30K non-cortical voxels) were demeaned and filtered (0.01 - 0.1Hz). Gray matter (GM), white matter (WM) and cerebrospinal fluid (CSF) signals were regressed, which is equivalent to global signal regression (GSR). The spatial dimension was reduced to 360 cortical parcels (Glasser et al., 2016), and each parcel’s timeseries was normalized to zero mean and unit standard deviation.

### Main algorithm to detect a QPP

The original method to detect a QPP, developed by Majeed and colleagues (Majeed et al., 2011), is centered around an algorithm that identifies similar segments of a rsfMRI scan using an iterative, correlation-based approach, then averages these segments to create a representative spatiotemporal template (Fig.S2). In this algorithm, a segment with a preset duration (here, ~20s or 30 consecutive timepoints) is initially selected. This initial segment is flattened and correlated with all the segments of the scan, which are selected and flattened one after another in a sliding fashion. This results in a timecourse of correlation between the initial segment and the rsfMRI scan, where timepoints corresponding to local maxima above a preset threshold (referred to as maxima for brevity) are then identified. The segments of the scan starting at those maxima are similar to the initial segment and are averaged together. The process is then repeated with the average of segments in place of the initial segment until negligible change between iterations is reached. The outputs of the algorithm are the final spatiotemporal QPP template obtained by averaging similar segments, and the correlation timecourse that shows the correlation of the final template with the scan at each point in time. The maxima in the correlation timecourse indicate the start of the segments that contribute to the final template, also referred to as the timepoints when the final template occurs.

### Obtaining group QPP1: an overview

The original method to obtain the group QPP1 (Majeed et al., 2011) involved concatenating the rsfMRI scans of individuals, running the described algorithm for a limited number of randomly selected initial segments, and from the resulting final templates, selecting the most similar one to the others as the QPP. We implemented a number of modifications to the original method, described in the sections that follow.

### Robust detection of QPP1 in individuals

To obtain group QPP1, we built upon our recent method improvements (Yousefi et al., 2018) by first robustly detecting QPP1 of individuals (Fig.S3). After concatenating the four preprocessed scans of each individual, *all possible initial segments* were examined using the main QPP algorithm. For each resulting template, the sum of correlation values at maxima was found. QPP1 was selected as the template with the maximum sum, which reflects a combination of high strength and frequent occurrence. In humans, QPP1 is ~20s long (Majeed et al., 2011; Yousefi et al., 2018; Abbas et al., 2019a,b; Briend et al., 2020); hence, we preset the segment duration to 30 timepoints (21.6s). In the prior studies (Majeed et al., 2011; Yousefi et al., 2018; Abbas et al., 2019a, 2019b; Belloy et al., 2018a, 2018b) the correlation threshold was set to 0.1 for the first three iterations of the main algorithm, and 0.2 for the remaining iterations; however, we increased this threshold to 0.2 for the first two iterations and 0.3 for the remaining iterations, mainly because of high quality and large dataset used here.

### Phase-adjusting individual QPP1s

A QPP is a spatiotemporal template that involves a cycle of activation and deactivation of different areas with different phases, and the detected QPP of each individual can be at any phase of the cycle in a given area. Proper averaging of the QPPs across individuals to move to the group level requires each QPP to have a certain phase in a reference parcel (referred to as the seed parcel for phase-adjustment; here chosen as left early visual cortex (V2)). For example (Fig.S4a), in an ideal phase-adjusted QPP, the 30-timepoint timecourse in V2 starts around zero at timepoint 1 and reaches its maximum before timepoint 15. To phase-adjust QPP1 of individuals, we used a roughly similar procedure to (Yousefi et al., 2018) that involves comparison of each QPP1 with all other templates, corresponding to all the examined initial segments. Out of the similar templates to QPP1, the one that met the criteria for an ideal phase was selected (see Fig.S4b for detailed description). Note that when comparing QPP1 with any other template, to account for minor differences in their phases, we used a fine phase-matching procedure, which involves shifting QPP1 a few timepoints forward and backward, and taking the maximum correlation across different time-shifts (see Fig.S5 for detailed description).

### Fine-tuned averaging of individual QPP1s

Before averaging the phase-adjusted QPP1 from all individuals, we included a fine phase-matching stage to mitigate the imperfections of the phase-adjustment procedure (see Fig.S6 for detailed description). We then used the template resulting from averaging the individual QPP1s as a prior for group QPP1. We correlated this prior template with all the scans of all individuals, identified the supra-threshold local maxima in the correlation timecourse, and used those timepoints as the start of the contributing segments to the final group QPP1. This last stage further mitigated any imperfections in the phase-adjustment procedure and ensured that group QPP1 (hereon, referred to as QPP1 for brevity) is simply an average of similar segments across the individuals.

### QPP1 in grayordinates

The reduction of the spatial dimensions from grayordinates to cortical parcels is a very effective and practical step that facilitates identification of the segments that contribute to QPP1. QPP1 in grayordinates can then be constructed by averaging the same contributing segments over the grayordinate timeseries instead of the parcellated timeseries (Fig.S1b). Unlike prior work, we did not place a threshold on the amplitude of QPP1 when visualizing in grayordinates, making it possible to observe more subtle trends of activation and deactivation. For all analysis beyond qualitative visualization, however, only vertices/voxels that exhibit statistically significant activation or deactivation at some time interval during the 20 s duration of the QPP1 were included. Statistically significant activity was defined as surpassing the 99^th^ percentile of a null distribution of activity created by averaging randomly selected spatiotemporal segments (for details, see Supplemental Material: Statistical evaluation of activity within QPPs, hereon referred to as S.M. for brevity, parts 1.1,1.2; also see Fig.S7-8).

### Summarizing the activity within QPP1 as a basis for further statistical analyses

To obtain a coarse summary of the activity within QPP1 in grayordinates, we clustered its timecourses, by comparing each pair of timecourses with the fine phase-matching procedure (maximum timeshift: ±2 timepoints). The upper triangle of the comparison matrix was then used to build the distance vector for the hierarchical clustering. We set the cut-off to 0.1 and kept the first ten largest clusters (These values, although somewhat arbitrary, have negligible influence on the results and do not change our conclusions). The two largest clusters are prominently larger than the others. One has the most spatial overlap with the cortical nodes of the default mode network (DMN), and we designated it as the first cluster (appearing first in all figures and tables). The other large cluster has timecourses which are strongly anticorrelated with those of the first cluster, and we designated it as the second cluster. Other clusters of QPP1 were sorted based on their median times of peak activation and deactivation, out of which, the clusters with intermediate timing and location relative to the first two clusters are referred to as the transitory clusters. To obtain a more detailed summary of the activity within QPP1, for each timecourse of QPP1, we also found the times of peak activation and deactivation (the latter as supplementary results; also called time of dip). For these summary maps, which are also the basis for the further statistical evaluations or comparisons, we only included the timecourses that exhibit statistically significant activity.

### Evaluating the coordinated propagation of activity within QPP1

To evaluate the coordinated propagation of of activity within QPP1, t-tests between pairs of clusters of timecourses, corrected for multiple comparisons, were used to determine significant differences in the time of peak/dip (see S.M.2 for details). The progression of times of peak/dip across clusters of timecourses, taken together with the spatial order of the clusters, demonstrates propagation of activity. Moreover, time-locked activity (coactivity) between non-adjacent areas across the whole brain is demonstrated by those non-adjacent areas being part of the same cluster.

### Existing parcellations, functional networks and gradients

Activity within QPP1, as a simple average of similar segments of rsfMRI timeseries, can be described and evaluated at the level of the whole brain, without the need for division into anatomical regions, networks or parcels. However, for multiple purposes, one being to compare QPP1 with the existing resting-state networks (RSNs) and functional connectivity gradients (FCGs), we adopted the parcellation and gradient schemes listed in Table S1 for seven brain regions of cerebral cortex, cerebellum, thalamus, hippocampus, amygdala, brainstem and deep brain nuclei, and striatum (also see Fig.S9). To quantify such comparison (see S.M.3 for details), first, we found the correlation between QPP1’s time of peak map and cortical FCG1 (Margulies et al., 2016); strong correlation supports propagation of activity within QPP1 is also consistent with FCG1. Although we only qualitatively compared QPP1 with the existing non-cortical FCGs, the statistically evaluated summary maps of QPP1 per each region were the basis for such comparison. Next, we grouped QPP1’s timecourses into the cortical RSNs (Yeo et al., 2011) and performed t-tests between pairs of groups, corrected for multiple comparisons, to determine the significance of differences in the time of peak activation; shifts of a few timepoints between pairs of RSNs support our statements about the sequential activation of RSNs within QPP1. Finally, we found the size of each cluster of QPP1’s timecourses per network/parcel per brain region, to show the extent of correspondance between QPP1 and particularly the non-cortical parcellation schemes.

The adopted parcellations were also used to identify different brain areas, as well as to describe the activity within QPP1 using established terminology. For example, we refer to the cortical areas that belong to the first cluster as the cortical nodes of the DMN, because of the extensive spatial overlap described earlier. As another example, we refer to the cerebellar areas that belong to the first cluster as the cerebellar areas that coactivate with the cortical nodes of the DMN.

### Definition of new terms

To simplify the description of the activity within a QPP, we define two new terms, again utilizing the adopted parcellation and gradient schemes. *(I) Propagation axis*. Within a QPP, simultaneous and consistent propagation of activity from all the nodes that constitute a RSN (e.g., RSN1) to all the nodes that constitute another RSN (e.g., RSN2) often occurs. We state that the activity propagates from the RSN1 to the RSN2 or that the propagation axis is RSN1→RSN2. When an existing FCG maximally separates RSNs 1 and 2, we also state that the activity propagates along that FCG. *(II) LPCC switching timepoints*. As a reference for describing the timing of activity within a QPP, we built a timecourse by averaging all the QPP’s timecourses that belong to the five central parcels of the left posterior cingulate cortex (LPCC) (Glasser et al., 2016). We define the coarse range of timepoints that the LPCC switches from deactivated to activated, or vice versa, as the LPCC switching timepoints. Any parcel could have been chosen for reference to describe the timing of activity within QPPs (for example, left V2, which was used as the seed parcel for phase-adjustment). We chose the LPCC because it is a prominent node of the DMN and the DMN is a prominent network in QPP1.

### Timing diffferences between brain regions

To examine the nuanced timing differences between brain regions that can suggest the driving mechanisms between them, per each brain regions, we found the number of vertices/voxels with a significant peak at each timepoint of QPP1, resulting in a histogram with 30 bins corresponding to the timepoints of QPP1. These historgrams are mostly bimodal, because of two the large anticorrelated clusters, and for the scope of this work, we only focused on the second mode, identified as the entries above the mid point of the cortical distribution. Only including the times of peak later than such midpoint, we used t-tests between pairs of regions, corrected for multiple comparisons, to determine the significant differences (see S.M.4 for details).

### Detection of additional QPPs

To examine whether additional recurring spatiotemporal patterns of activity are present in the rsfMRI timeseries, we regressed QPP1 and reanalyzed the residuals. Two methods for regression were implemented, both using GLM, and performed at the individual level for each of the four scans. First, scan-wise regression (Fig.S10a), where QPP1 was convolved with its correlation timecourse to build the regressor. Since QPP1 is a spatiotemporal pattern and its timecourse are different for each parcel, the timecourse for each parcel was convolved with the QPP1’s correlation timecourse and the result was regressed from that parcel’s timeseries. Second, segment-wise regression (Fig.S10b), where QPP1 was regressed from each of its contributing segments, which were replaced by the residuals. To ensure no similar segments to QPP1 exist in this residual scan, QPP1 was correlated with the residual scan and was regressed from segments corresponding to any detected supra-threshold local maxima. The two regression methods have similar outcomes (Fig.S11a), and we based our group-level report on the scan-wise regression, which runs faster.

After QPP1 of each individual was regressed per scan, each parcel’s timeseries was normalized to zero mean and unit standard deviation and concatenated across the four scans. The secondary QPP (QPP2) was then detected and phase-adjusted following the same methods used for QPP1. We further regressed QPP1 and QPP2 of individuals and reanalyzed the residuals to detect and phase-adjust QPP3. We limited the scope of this work to QPPs 1 to 3. When building QPP2 and QPP3, averaging the contributing segments over the original scans or the residual scans resulted in nearly identical templates (Fig.S11b). We therefore averaged the segments from the original scans to avoid possible minor distortions induced by regression. Group QPPs 2 and 3 were obtained for the parcellated data and then for grayordinates, summarized and statistically evaluated using the same methods as for group QPP1.

### QPP basic metrics and transition count

For each QPP, we can readily find its strength (the median of the supra-threshold local maxima in the QPP’s correlation timecourse with the scan) and its occurrence interval (the median of the time interval between successive maxima in the correlation timecourse). These metrics reflect how well each QPP represents its contributing segments, and how often it occurs. The strength and occurrence interval of QPPs 1-3 were calculated at the individual level and averaged across individuals. We also characterized transitions between QPPs, by counting the number of times that a contributing segment of QPPi was followed by a contributing segment of QPPj. This resulted in a 3×3 matrix for each individual, which was summed across individuals to obtain the group level matrix.

### Contribution to functional connectivity

To examine the extent to which patterns of activity represented by the QPPs contribute to functional connectivity, we calculated the Pearson correlation between each pair of the 360 cortical areas using the original timeseries and the residual timeseries after regressing QPPs 1-3, using the scan-wise method. For this calculation, we first sorted the 360 cortical parcels based on the seven RSNs (Yeo et al., 2011), using a simplified rule to assign each parcel to the RSN with which it has maximal overlap. The resulting functional connectivity matrices were averaged across individuals after Fisher transformation. To characterize these matrices at a high level, we created histograms of correlation values for all pairs of areas for four cases: before regression of any QPPs, after regression of QPP1, after regression of QPPs1 and 2, and after regression of QPPs 1-3. We then determined the percentage of correlation values above 0.1 or below −0.1 for each case (0.1 was chosen somewhat arbitrarily to be qualitatively meaningful but is above the 99^th^ percentile of null values built by phase-shuffling the timeseries and applying the abovementioned procedure). We further performed the following complementary analyses. First, the variance of the functional connectivity matrix before and after regression of each QPP was calculated. To obtain null values, the QPPs were convolved with their shuffled correlation timecourses, regressed scan-wise, and the variance of the functional connectivity matrix was found. Second, the correlation between the functional connectivity matrix before regression of any QPP and the functional connectivity matrix after regression of each QPP was calculated. Finally, the correlation between 360 cortical areas within each QPP (i.e., functional connectivity within QPP, over the ~20 s timecourses) was calculated and qualitatively compared with the functional connectivity matrices before and after regression of that QPP.

### Robustness analyses

In our preprocessing pipeline, GM was regressed along with WM and CSF (i.e., GSR). Although GSR is a controversial practice (Power et al., 2017), our recent work indicates that it improves the correspondence of QPPs across individuals (Yousefi et al., 2018). To examine the influence of GSR on our results, we averaged the contributing segments of each QPP over the WM and CSF regressed timeseries and compared the resulting template with that QPP (obtained after GSR) at the individual level. To further examine the effects of any nuisance regression or filtering on the main features of the QPPs at the group level in grayordinates, we averaged the contributing segments of each QPP over timeseries that were only demeaned and visually compared the resulting template to that QPP (obtained after filtering and nuisance regression). To examine the effect of choosing a particular area as the seed for phase-adjustment, for each group QPP, we chose another seed parcel to obtain a template with a reversed phase to that QPP and visually compared the results (LPCC was used for QPP1, left supramarginal gyrus (smg) for QPP2, and left primary motor area (M1) for QPP3). To examine the effect of correlation threshold, we compared group QPPs of forty randomly selected individuals detected based on the setting here (0.2 and 0.3) versus the setting in the prior work (0.1 and 0.2). To test the reproducibility of the QPPs, individuals were randomly divided into two equal subgroups, 50 times, QPPs 1-3 were obtained for each subgroup, and compared across subgroups.

All codes of this work is openly available at https://github.com/GT-EmoryMINDlab and QPP analysis is also implemented in the C-PAC pipeline (www.nitrc.org).

## Results

As previously reported, QPP1 involves a cycle of activation and deactivation of different brain areas with different relative timings and includes propagation of activity in the medial prefrontal cortex (Fig.1). New to this study, the cycle of activity and propagation extends throughout and beyond the cortex to the entire brain, including the cerebellum, thalamus, striatum, hippocampus, amygdala and brainstem. An in-depth description of the most prominent features is as follows.

**Figure 1.**
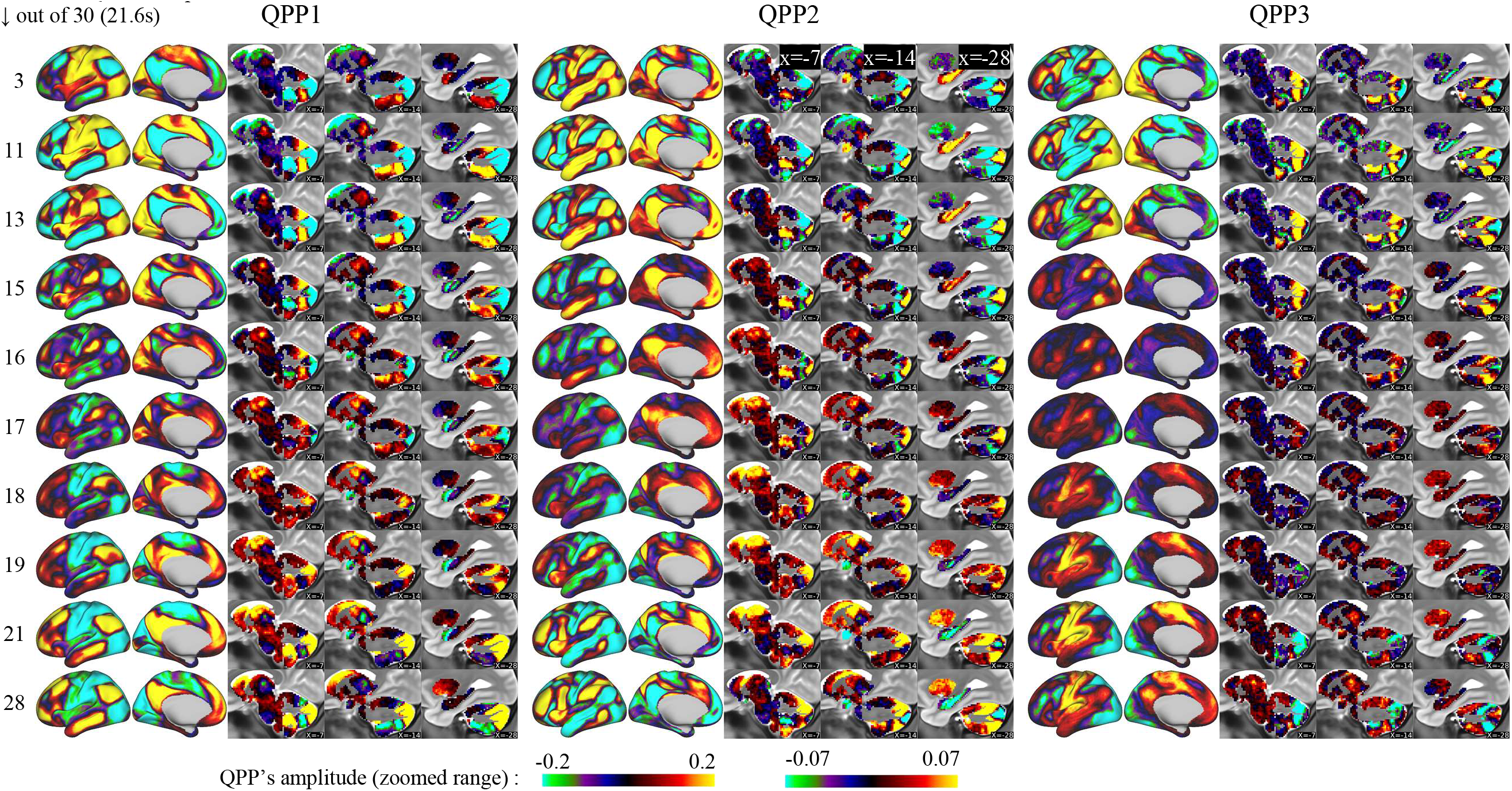
QPPs 1-3 involve coordinated propagation of activity along functional connectivity gradients (FCGs) across the whole brain. Each row corresponds to a timepoint of the QPP and shows the level of activation or deactivation of brain areas with warm and cool color ranges, respectively. For all 30 timepoints see Videos 1.

### Propagation of activity along cortical FCG1

Within QPP1 (Fig.1, Video 1, Table 1), activity propagates along the primary cortical FCG (FCG1), from areas that belong to the somatomotor network (SMN) to all nodes of the DMN. Deactivation begins in the lower and upper limb areas of the SMN and spatially expands to the supplemental motor area and the surrounding nodes of the dorsal attention network (DAN), such as the premotor area and the superior parietal lobe (Fig.1 timepoints 11-16). Activation expands from each node of the ventral attention network (VAN) to a neighboring node of, first, the frontoparietal network (FPN), and later, the DMN (Fig.1 timepoints 16-19). For example, activity expands from the anterior cingulate cortex (ACC) to the ventromedial prefrontal cortex (vmPFC) (see Table 2 for the list of all nodes). Clusters of QPP1’s timecourses, with their spatial order (Fig.2a,b) and significant progression of timing (Fig.S12a-d, Cortex), together with the times of peak activity of QPP1’s timecourses (Fig.2c), which is strongly correlated with FCG1 (Fig.S13a), summarize the propagation along the cortical FCG1 (also see Fig.S13c-e, S14a that support the sequential activity of RSNs). Although the FPN is known to be part of the TPN (Petersen and Posner, 2012), within QPP1, it is positively correlated with the DMN and anticorrelated with the DAN and VAN (Fig.S13d). Moreover, the visual network (VN) is anticorrelated with the FPN/DMN. The primary visual area (V1), however, exhibits clear propagation of activity from the peripheral areas to the foveal areas, with the foveal areas being positively correlated with the FPN/DMN (Fig.S15a-b). Such focal propagation is qualitatively along the V1’s FCG1 (Haak et al., 2018) and in line with Hindriks et al., (2019).

**Figure 2.**
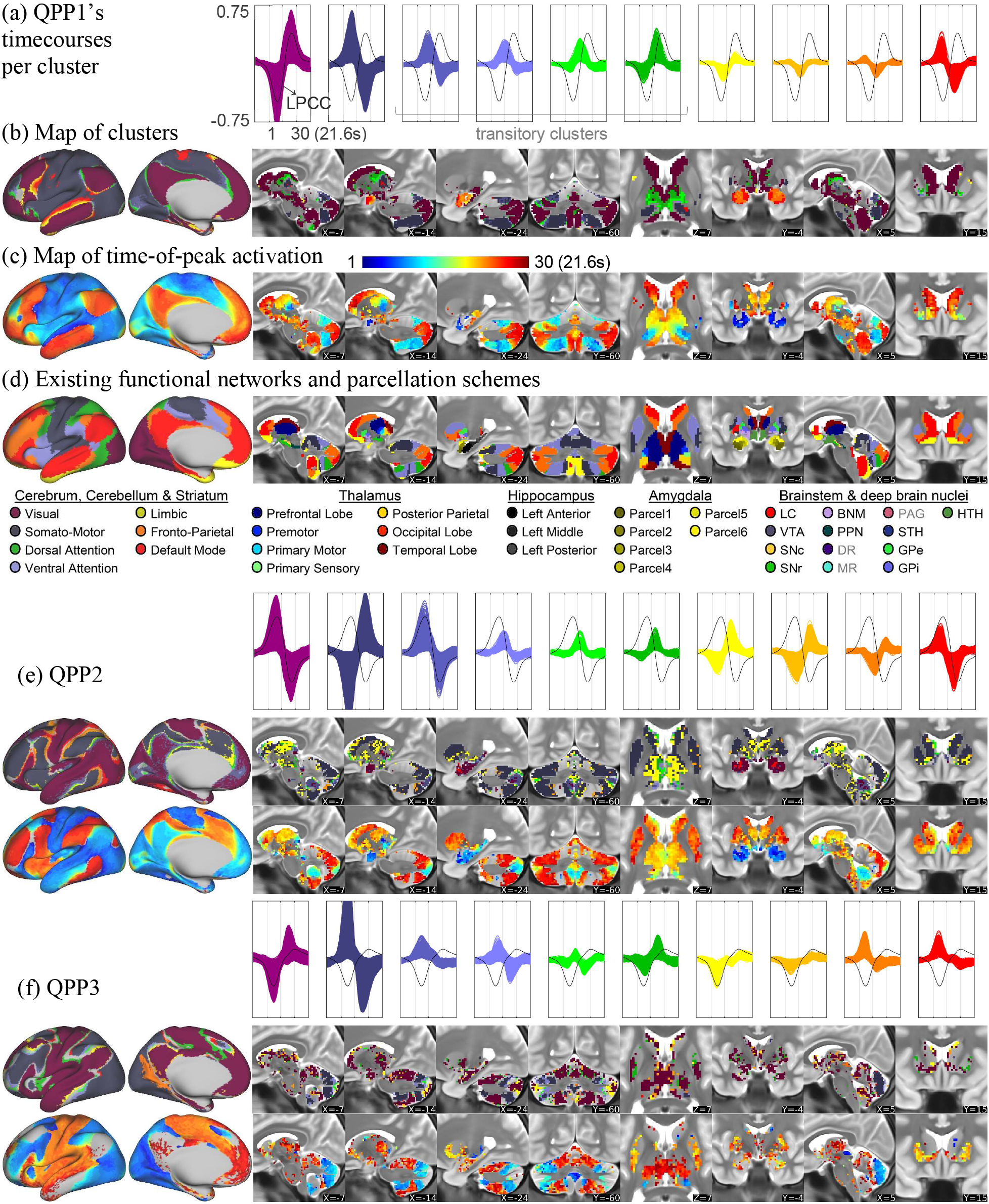
Summary maps of activity within QPPs, illustrating coordinated propagation across the whole brain. (a) QPP1’s timecourses (plotted together) per cluster (LPCC timecourse in black for reference). (b) Brain areas, at the left cortical hemisphere and representative non-cortical planes, color-coded based on correspondence with QPP1’s clusters. (c) Timepoints of peak activation of QPP1’s timecourses. (d) Existing functional networks and parcellations for comparison. QPPs 2 (e) and 3 (f).

**Table 1.** Axis of propagation of activity within QPPs

|  |  |  |
| --- | --- | --- |
| Q<br>P<br>P<br>1 | Cerebral cortex | Cortical FCG1, SM→TP→DM<br>Finer axis: SM→DA & VA→FP/DM & V↓↑FP/DM |
|  | Cerebellum | Cerebellar FCG1, DA/VA→FP/DM |
|  | Thalamus | Novel, posterolateral → medial (or SM→FP/DM) |
|  | Hippocampus | Hippocampal FCG1, Posterior → Anterior |
|  | Amygdala | Novel, Center → Sides |
|  | Brainstem | FP/DM |
|  | Striatum | Striatal FCG, Caudal/Orbital → Rostral & Lateral → Medial |
| Q<br>P<br>P<br>2 | Cerebral cortex | Cortical FCG3, TP→SM/V & DM→TP<br>Finer axis: DA→SM/V & DM→FP→VA→DA |
|  | Cerebellum | Cerebellar FCG2, DM→FP/VA/DA |
|  | Thalamus | Novel, most medial → anterolateral (or DM→FP/VA) |
|  | Hippocampus | Hippocampal FCG1, Anterior → Posterior |
|  | Amygdala | SM/DM |
|  | Brainstem | FP/VA |
|  | Striatum | Striatal FCG, Rostral → Caudal & Medial → Lateral |
| Q<br>P<br>P<br>3 | Cerebral cortex | Cortical FCG2, V↓↑SM, also V/TP→SM/DM<br>Finer: V/DA/FP↓↑VA/SM/DM, also V/DA/FP→VA→SM→DM |
|  | Cerebellum | DA/FP↓↑SM/DM, also DA/FP→DM |
|  | Thalamus | V↓↑ SM/DM |
|  | Hippocampus | SM/DM |
|  | Amygdala | SM/DM |
|  | Brainstem | SM/DM (particularly, pons) |
|  | Striatum | SM/DM (particularly, posterior putamen) |

Cortical networks of SM (somatomotor), TP (task positive), DM (default mode), DA (dorsal attention), VA (ventral attention), FP (frontoparietal), V (visual) and L (limbic) ↓↑ (anticorrelation), single entry without → (simple coactivity)

**Table 2.** Approximate cortical nodes that constitute the networks in the finer axes of propagation in Table 1

|  |  |  |
| --- | --- | --- |
| Q<br>P<br>P<br>1 | SM→DA | limb/face → preM/SPL<br>foot/genitalia → sma |
|  | VA→FP/DM | ACC → vmPFC<br>aI → ifg<br>ifs → sfg/mfg/ifg<br>mTL: posterior → anterior<br>Ag/PCC: surround → center |
|  | DM→LNG | TL: medial → superior<br>left 55b/IFJ/SFL: anterior → posterior |
| Q<br>P<br>P<br>2 | DA→SM/V | DA-bordering-SM/V→SM/V |
|  | DM→FP→<br>VA→DA | vmPFC → aACC → dACC<br>ifg → aI → mI → inferior-preM<br>sfg/mfg/FPC → ifs<br>TL: anterior→ infero-posterior →posterior<br>Ag → smg → PF/SPL<br>PCC→RSC/POS → postero-medial-VA/DA |
| Q<br>P<br>P<br>3 | V/DA/FP→<br>VA→SM | TPOJ → inferior-TPJ → inferior-PF → face<br>aI → mI → inferior-preM → face<br>sfg → FEF → limb<br>SPL/sma → limb/foot |

preM (premotor area), SPL (superior parietal lobe), sma (supplementary motor area, however, NOT a node of Yeo's DAN), ACC (anterior cingulate cortex), vmPFC (ventromedial prefrontal cortex), aI (anterior insula), ifg (inferior frontal gyrus), ifs (intermediate frontal sulcus), sfg (superior frontal gyrus), mfg (middle frontal gyrus), mTL (middle temporal lobe), Ag (angular gyrus), PCC (posterior cingulate cortex), LNG (language areas), TL (temporal lobe), 55b (area 55b), IFJ (inferior frontal junction), SFL (superior frontal language area), aACC (anterior ACC), dACC (dorsal ACC), mI (mid insula), FPC (frontopolar cortex), smg (supramarginal gyrus), PF (area PF), RSC (retrosplenial cortex), POS (parieto-occipital sulcus), TPOJ (temporo-parieto-occipital junction), TPJ (temporo-parietal junction), FEF (frontal eye field).

### Propagation of activity in other brain regions

Within QPP1, in addition to the cerebral cortex, most other regions of the brain exhibit propagation of activity, time-locked to the cortical propagation, in directions that are consistent with the cortical propagation and which lie along all FCGs reported so far. Summary maps of activity generally match the adopted parcellation schemes and the known cortico-subcortical tract-based connections. A detailed description for each region follows (for each region, see Fig.2a-d as the spatial reference, Fig.S12a-d and Table S2 as the quantitative support)

*Cerebellar* activity generally matches the adopted parcellation scheme (Buckner et al., 2011), in that the cerebellar areas recognized as DMN, for instance, in that scheme are strongly correlated with the cortical nodes of the DMN within QPP1. As in the cortex, cerebellar areas that coactivate with the cortical FPN and DMN are positively correlated with each other and are anticorrelated with the cerebellar areas that coactivate with the cortical DAN/VAN. At the LPCC switching timepoints, activation expands from areas that coactivate with the DAN/VAN to areas that coactivate with the FPN/DMN. Such propagation is qualitatively along the cerebellar FCG1 (Guell et al., 2018).

*Thalamic* areas that are coactive with the cortical areas of the SMN and DAN are mainly located posterolaterally, in line with their tract-based connections (Behrens et al., 2003). For example, the most posterolateral part of the thalamus (possibly the foot area of the ventral posterior nucleus (Sherman and Guillery, 2013)) is coactive with the medially located cortical foot area. As in the cortex, activity expands from areas coactive with the SMN to areas coactive with the FPN/DMN. Given the synchrony and spatial consistency of thalamic and cortical activity and the finding that cortical activity sweeps FCG1 during QPP1, the propagation of activity across the thalamus suggests a previously-undescribed macroscale FCG for this region.

The posterior part of the *hippocampus* coactivates with the cortical nodes of the DMN, while its anterior part coactivates with the amygdala. This is in accord with the consensus that the hippocampus is a node of the DMN and exhibits functional specialization along its long axis, with its anterior part closely interacting with the amygdala (Robinson et al., 2016). Activity propagates along the hippocampus from posterior to anterior, qualitatively sweeping the hippocampal FCG1 (Vos De Wael et al., 2018).

Propagation of activity towards the *amygdala* via the hippocampus happens at the same time that activity propagates from the medial temporal lobe (TL) towards the superior TL (Fig.S15a,c). Deactivation propagates across the amygdala from the center to the sides, qualitatively sweeping across the main parcels (Tyszka and Pauli, 2016), suggesting a FCG across the amygdala that matches a focal gradient across the cerebral cortex.

*Brainstem and deep brain nuclei* primarily coactivate with the cortical nodes of the FPN/DMN. While difficult to resolve, the probable locations of dopaminergic substantial nigra pars compacta (SNc) contains the most voxels with the earliest peak times, with the majority of the earliest voxels not belonging to any adopted map of nuclei (note Fig.S12d versus Table S2 also Fig.S16). Voxels which are anticorrelated with the FPN/DMN seem to be located in the pons, possibly in the pontine nucleus that relays the cortical inputs to the cerebellum.

*Striatal* areas are mostly coactive with the cortical nodes of the FPN/DMN. The ventrolateral striatum and the tail of the caudate (for the latter, note Fig.S16), however, are coactive with the cortical transitory clusters. Activation expands from areas that are coactive with the cortical transitory areas to the areas that coactivate with the cortical FPN/DMN. This propagation is qualitatively along the striatal FCG (Marquand et al., 2017) and matches the topographical cortico-striatal tract-based connections (Choi et al., 2012; Marquand et al., 2017).

### Timing differences across brain regions

As activity propagates in most brain regions within QPP1, the numbers of vertices/voxels that peak at each timepoint of QPP1 in each region (Fig.3a) suggest driving mechanisms. The cerebellum slightly but significantly lags the cortex (see Fig.S17 for statistical support throughout this part), and such lag is in line with Marek et al., (2018). In contrast, as activity propagates to the cortical nodes of the FPN/DMN, the coactive thalamic areas lead by a median of 4 timepoints (2.9s). Brainstem and deep brain nuclei also peak earlier than the cortical nodes of the FPN/DMN, leading with a median peak time of 2 timepoints (1.4s). Interestingly, timing differences between the abovementioned regions remain similar if only the vertices/voxels that belong to the first cluster of QPP1 are considered (Fig.S12a-c), where, for example, the thalamus leads the cortex by 3 timepoints.

**Figure 3.**
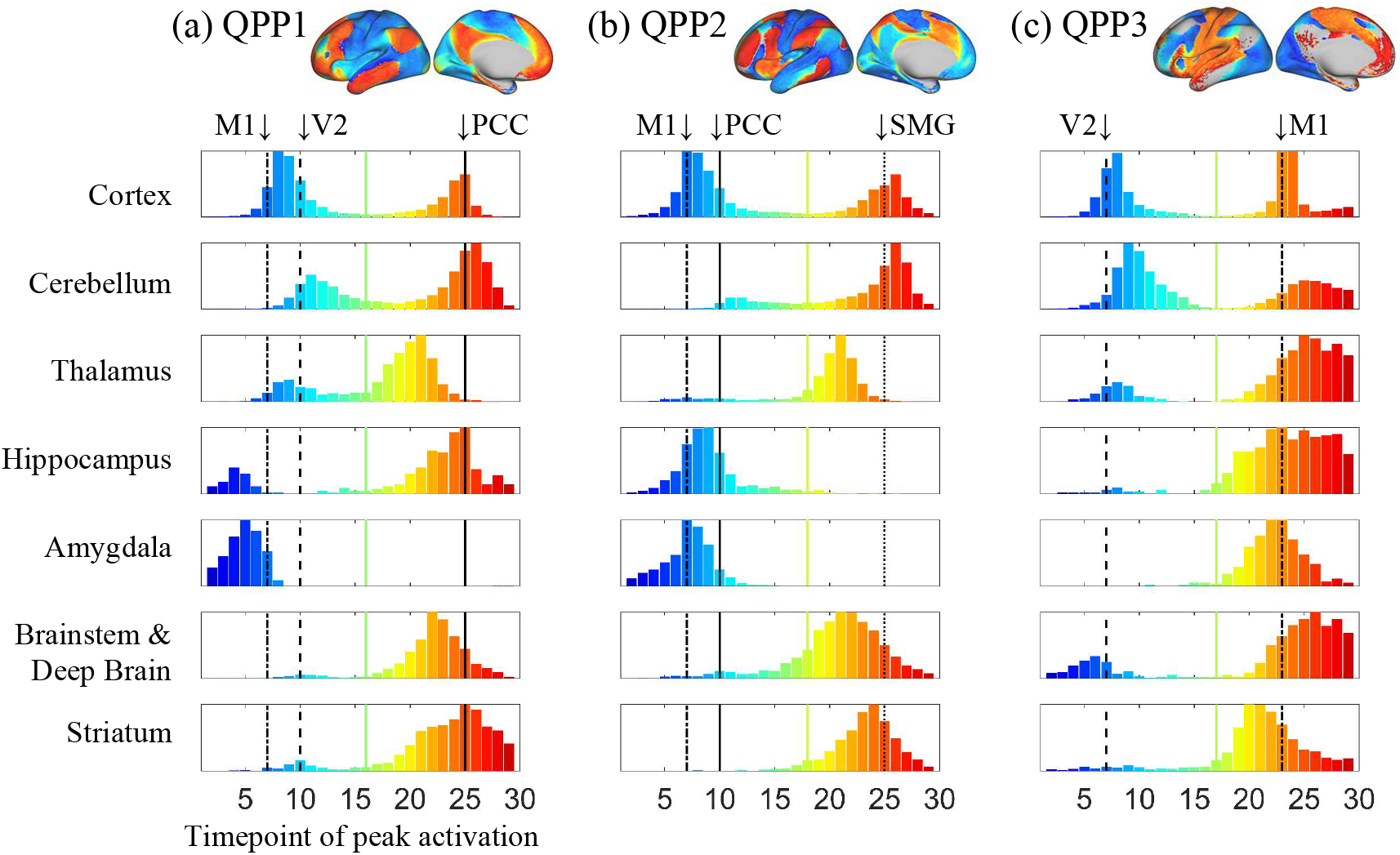
The number of vertices/voxels with peak activity at each timepoint of QPPs 1-3 at each brain region. As qualitative references, the times of peak activity at the cortical areas of the left PCC, V2, M1 and SMG are plotted in black with unique line types. The distributions are mostly bimodal and the comparison of time difference between pairs of regions were done on the second mode, identified as the entries above the mid timepoint of the cortical distribution (indicated by the colored line for each QPP). Thalamic and brainstem and deep brain areas lead the cortex in QPPs 1 and 2 as activity propagates to the cortical nodes of FPN/DMN in QPP1 and the cortical nodes of FPN/VAN in QPP2. The cerebellum slightly lags the cortex in all QPPs.

### Propagation of activity along other FCGs

QPPs 2 and 3 have distinct spatiotemporal characteristics (compared to QPP1 and to each other) but still involve a coordinated propagation of activity throughout the whole brain. The direction of propagation is consistent with existing FCGs and generally matches the adopted parcellation schemes and the known tract-based connections (for the following results, refer to Fig.1, Fig.2e-f, Video 1 and Table 1–2, and for the statistical support see Fig.S18-23 and Table S3-4).

*Within QPP2*, activity propagates along the cortical FCG3. Deactivation initially starts in the areas of the DAN that lie in the border with the SMN and VN, and from there deactivation spatially expands to the SMN and VN. Furthermore, activity expands from nodes of the DMN to neighboring nodes of the FPN, then the VAN, and finally to the DAN. As in QPP1, V1 exhibits distinct activity compared to other areas of the VN, with the foveal area now in phase with the FPN/VAN.

Within QPP2, *cerebellar* areas labelled as DMN in the adopted parcellation scheme are strongly correlated with the cortical nodes of the DMN and anticorrelated with the cerebellar areas labeled as DAN, VAN, and FPN (or collectively TPN). As in the cortex, at the LPCC switching timepoints, activation expands from areas that coactivate with the cortical DMN to areas that coactivate with the cortical TPN, which is also qualitatively consistent with the cerebellar FCG2 (Guell et al., 2018).

*Thalamic* areas coactive with the cortical SMN and VN are mainly located posterolaterally. The majority of the remaining thalamus is coactive with the cortical FPN/VAN, a smaller medial portion is coactive with the cortical transitory clusters (particularly note Fig.S20), and a very small medially-located portion, likely in the mediodorsal nucleus (MD), is coactive with the cortical DMN. Consistent with the cortex, at the LPCC switching timepoints, activity expands from the medial-MD to the lateral-MD and further to the antero-lateral thalamus. This may be evidence for the existence of another macroscale FCG across the thalamus.

*Hippocampus* and *amygdala* are coactive with the cortical DMN, yet peak slightly earlier (particularly the amygdala; note Fig.S20). Furthermore, the posterior part of the hippocampus slightly lags the anterior, which indicates propagation along the long axis, or qualitatively along the hippocampal FCG1, but now from anterior to posterior.

*Brainstem and deep brain nuclei* now primarily coactivate with the cortical nodes of the FPN/VAN and the relative number of the early voxels compared to FPN/VAN has increased (compare Fig.S18d with S12d, also note Fig.S20).

*Striatal* areas are mostly coactive with the cortical nodes of the FPN/VAN. As in the cortex and most other non-cortical regions, at the LPCC switching timepoints, activity expands from the DMN-coactive to the FPN/VAN-coactive areas, along both rostro-caudal and medial-lateral directions, which are also qualitatively along the striatal FCGs.

### Timing differences between regions within QPP2

As in QPP1, the cerebellum slightly but significantly lags the cortex (Fig.3b, S17). As activity propagates to the cortical nodes of the FPN/VAN, the thalamus leads by a median of 4 timepoints (2.9s), and the brainstem and deep brain nuclei lead by a median of 3 timepoints (2.1s). Unlike in QPP1, the thalamus and brainstem areas are now more comparable in timing. As in QPP1, timing differences between the abovementioned regions remain when only the vertices/voxels that belong to the second cluster of QPP2 are considered (Fig.S18a-c).

Activity within *QPP3* is consistent with the cortical FCG2. The cortical areas of the SMN and VN exhibit a simple cycle of activation and deactivation with an opposite phase relative to one another. Nodes of the VAN and DMN are correlated with the SMN and nodes of the DAN and FPN are correlated with the VN. As activation levels are switching in the areas of the VN and SMN and for a short time afterwards, focal propagations occur via the VAN to the SMN and later towards the DMN.

*Cerebellar* coactivity still generally matches the adopted parcellation scheme and exhibits propagation consistent with the cortex with a slight yet significant lag (Fig.3c, S17). Note the lobules I-IV, labeled as primary SMN in the adopted scheme, exhibit strong activity, not present in QPPs 1 and 2. Other than a small area in the posterolateral *thalamus* that is coactive with the cortical VN, the remaining thalamic areas, *hippocampus, amygdala*, posterior *putamen*, and *pons* (or possibly the pontine nucleus) are all mostly coactive with the cortical SMN/DMN. The striatum and amygdala lead cortical SMN/DMN by a median of 2 timepoints (1.4s).

### QPPs basic metrics and transition counts

QPPs 2 and 3 are comparable to QPP1 in strength, but they occur less often than QPP1 (Table 3). Each of QPPs 1-3 is more often followed by a QPP of a different type rather than by the same type (Table 4). The QPP transition count matrix reflects the finding that QPP1 occurs most often, as most transitions occur to or from QPP1. Each QPP exhibits a unique pattern of correlation between pairs of areas, i.e., each QPP exhibits unique functional connectivity (note Fig.S13c, S19c, S22c, also Fig.4c). The correlation between pairs of matrices of functional connectivity within each QPP is 0.07 for QPPs 1 and 2, 0.04 for QPPs 1 and 3, and 0.1 for QPPs 2 and 3.

**Figure 4.**
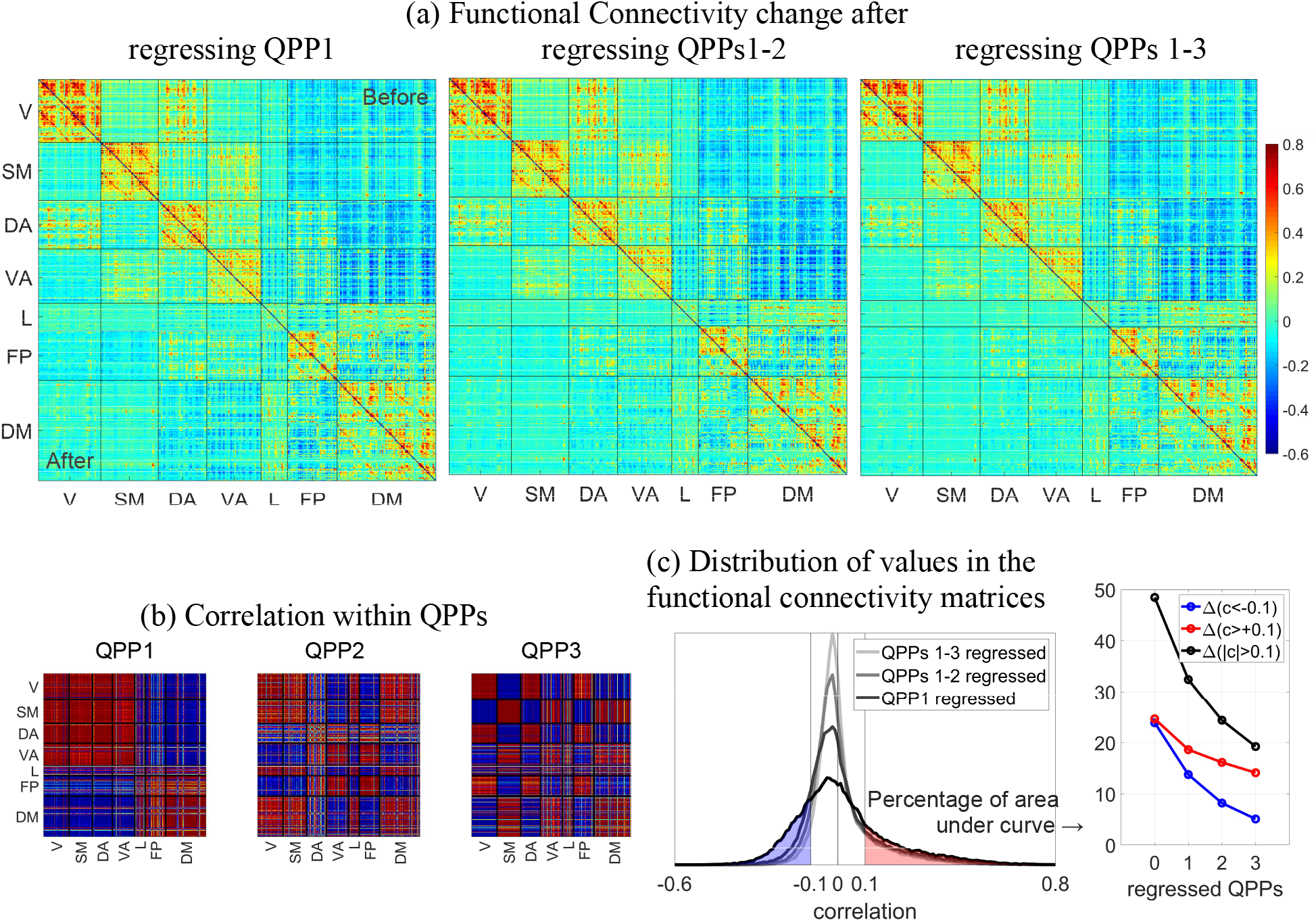
Change in functional connectivity between pairs of areas after regressing QPPs. (a) For each matrix, the top right half is the functional connectivity between 360 cortical parcels ordered based on seven RSNs, before regressing any QPP, and the bottom left half is the functional connectivity after scan-wise regression of each QPP. RSNs include networks of visual (V), somatomotor (SM), dorsal attention (DA), ventral attention (VA), limbic (L), frontoparietal (FP) and default mode (DM). Regressing each QPP progressively reduces the correlation between pairs of areas, within and particularly between RSNs. Such reduction is consistent with the correlation within each QPP, shown in (b). (c) The distribution of correlation values for each functional connectivity half-matrix shown in (a), and the percent of the correlation values above 0.1 or below −0.1 for each case. Both positive and negative correlation values are progressively reduced by regression of QPPs, but the negative correlation values are more affected.

**Table 3.** Basic metrics of QPPs (average± standard deviation)

|  | QPP1 | QPP2 | QPP3 |
| --- | --- | --- | --- |
| Strength | 0.40±0.021 | 0.37±0.02 | 0.35±0.02 |
| Occurrence interval (s) | 52.9±6.9 | 64.4±9.9 | 77.5±12.8 |

**Table 4.**
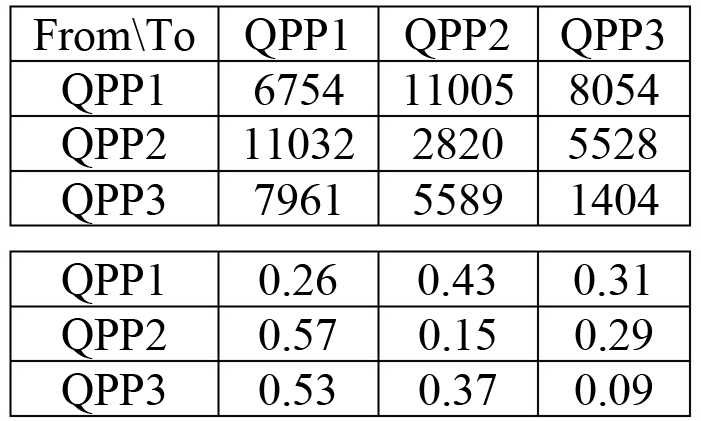
Transition count and probability of transition between QPPs

| From\To | QPP1 | QPP2 | QPP3 |
| --- | --- | --- | --- |
| QPP1 | 6754 | 11005 | 8054 |
| QPP2 | 11032 | 2820 | 5528 |
| QPP3 | 7961 | 5589 | 1404 |
| QPP1 | 0.26 | 0.43 | 0.31 |
| QPP2 | 0.57 | 0.15 | 0.29 |
| QPP3 | 0.53 | 0.37 | 0.09 |

### Contribution to functional connectivity

The strong recurring coordinated activity represented by QPPs contributes substantially to correlation between brain areas, which in turn is used to derive RSNs and FCGs. Regression of QPPs 1-3 from the rsfMRI timeseries progressively reduces the correlation between pairs of areas, within and particularly between RSNs (Fig.4a). In total, QPPs 1-3 account for ~61% (Fig.4c, based on start and end points in black) of the correlation values with a magnitude greater than 0.1. The number of pairs of areas with strong negative correlation (−1 to −0.1) is more drastically reduced by regression of QPPs (~79%, Fig.4c in blue) than the number of pairs of areas with strong positive correlation (0.1 to 1; ~43%, Fig.4c in red).

From the perspective of variance in the functional connectivity matrix, QPP1 explains ~37% of the original functional connectivity, QPPs 1-2 explain ~53%, and QPPs 1-3 explain ~63%. When shuffled correlation timecourses of QPPs are used, the variances of the resulting functional connectivity matrices after progressive regression of QPPs (corresponding to the lower triangles in Fig.4a) remain unchanged. After regression of QPP1, QPPs 1-2, or QPPs 1-3, the original functional connectivity matrix (upper triangles in Fig.4a) and the resulting functional connectivity matrices (lower triangles in Fig.4a) are uncorrelated (r<0.004). Within each RSN, particularly in the VN, SMN and DAN, functional connectivity between homologous parcels in the right and left hemispheres are the least affected by regressing QPPs 1-3 (note the parallel bands to the diagonal).

### Robustness analyses

When averaging the contributing segments of each QPP using the timeseries that were only WM and CSF regressed, we obtain patterns that are nearly identical to that QPP (Fig.S24), proving that GSR does not influence the reported QPPs. When averaging the contributing segments of group QPPs using the grayordinate timeseries that are only demeaned, but not filtered or nuisance-regressed, we obtain patterns that qualitatively have all the main characteristics reported here (Video 2), the most important being the coordinated propagation of activity across the whole brain; this proves the prominent features of QPPs are not artifacts of preprocessing. QPPs with the reversed phase (Video 3, Fig.S25, also see Fig.S26) exhibit all the main characteristics, proving robustness to the settings in the phase-adjustment. The slight change of the correlation threshold compared to previous works has no effect on the reported QPPs (Fig.S26). When individuals were randomly divided into two sub-groups, 50 times, the sub-group QPPs were nearly identical (medians across repetitions were 0.99, 0.98 and 0.97, for QPPs 1-3), proving QPPs are reproducible across different groups of individuals.

## Discussion

We have described three spatiotemporal patterns of activity (QPPs 1-3) that encompass the whole brain, propagate across the known macroscale functional connectivity gradients (FCGs), and account for most of the functional connectivity within and particularly between resting-state networks (RSNs). These patterns provide novel insights into the brain’s intrinsic activity that cannot be inferred by functional connectivity, RSNs or FCGs.

### Functional connectivity, RSNs and FCGs

Functional connectivity, typically calculated as the Pearson correlation between timeseries of different brain areas, describes the average relationships between areas over the course of an entire scan (e.g., ~10 minutes). Besides, time-lagged relationships can manifest as a lack of correlation or even anticorrelation between areas. In contrast, the spatiotemporal patterns of QPPs, which are the major contributors to functional connectivity, can describe nuanced time-lagged relationships that might vary between the same areas over the course of the scan.

Functional connectivity is the basis to derive RSNs and FCGs, both of which indicate spatial organization of the brain’s intrinsic activity with links to task-related activity and behavior. First few QPPs are major contributors to functional connectivity, involve sequential activity of RSNs and propagation of activity along FCGs. Therefore, very likely, first few QPPs also majorly contribute to RSNs and FCGs, the extent of which can be quantified by analyzing residuals after regressing the QPPs. Put in other words, RSNs and FCGs are likely snapshots of a few propagating patterns of coordinated activity that encompass the whole brain, from the brainstem to the cerebral cortex, and involve nuanced timing relationships between brain areas and regions. Note each QPP, that represents certain set of time intervals during the course of the rsfMRI scan, corresponds to a particular FCG, but the overlaid summary maps of QPPs correspond to RSNs.

A noteworthy insight provided by QPPs is about the widely documented intrinsic anticorrelated activity between the task positive network (TPN) and task negative or default mode network (DMN) (Fox and Raichle 2007). First, this intrinsic anticorrelation involves propagation of activity between the TPN and DMN, which might possibly be the case in the dynamics of task-related activity, similar to Kucyi et al., (2020). Second, the executive control or the frontoparietal network (FPN), which is a subnetwork of the TPN (Petersen and Posner, 2012), is in fact, positively correlated with the DMN at some time intervals (that contribute to QPP1) during the course of the rsfMRI scan.

Another noteworthy insight provided by QPPs is about the coactivation maps of the non-cortical regions with the cortical RSNs. For example, while the posterolaterally located unimodal thalamic areas coactivate with the unimodal cortical areas within all QPPs, a given voxel in other thalamic areas can coactivate with one cortical RSN (or the bordering areas between two RSNs) in one QPP and another cortical RSN (or the bordering areas between two other RSNs) in another QPP. Therefore, it might be a more robust approach to perform RSN-based parcellation in the thalamus using only certain time intervals of the rsfMRI scan and incorporate a few coexisting parcellation schemes (even consider coactivity with the bordering areas between RSNs), which is unlike the common approaches (Zhang et al., 2010; Ji et al. 2018). The same note holds for other non-cortical regions, such as the brainstem, striatum, amygdala and even the cerebellum.

### Timing differences between regions

QPPs provide suggestions about plausible driving mechanisms between brain regions. For example, the thalamus, brainstem and deep brain areas lead the cerebral cortex as activity propagates between the cortical nodes of the TPN and DMN in both QPP1 and QPP2, with the thalamus ~1.5 s and 0.7 s ahead of the brainstem. This implies a specific and key role for the thalamus in the intrinsic switching of the activity between areas involved in the externally oriented attention (TPN) and internally oriented attention (DMN), in line with the general role of the thalamus in the attentional control (Halassa and Kastner, 2017). It also suggests that the brainstem and deep brain nuclei, which promote arousal and reinforce attentiveness (Avery and Krichmar, 2017), join the thalamus in this key role.

### Comparison to other reports of propagation of activity

Propagation of activity is a prominent feature of the QPPs and has been reported in previous rsfMRI-based studies. Hindriks et al. (2019) reported propagation from the TPN to the DMN and simultaneously, from anterior to posterior V1, which are consistent with our results. Their argument that RSNs are likely to arise from propagating patterns is quantitatively illustrated here, as we show that three QPPs account for the vast majority of functional connectivity. Mitra and colleagues have reported propagation in terms of lag threads, which while intriguing, is complementary rather than comparable to our findings (Mitra et al., 2015, 2016). In their work, RSNs arise from one-directional motifs which are common among a few reproducible lag threads, no RSN leads or lags the other, and the range of lags in the threads are ~±1 second. In contrast, RSNs are readily observable within QPPs and they clearly exhibit lead/lag relationships at much longer time scales. To our knowledge, the extent of the coordination of propagation across the whole brain reported here and its relationship to the common descriptors of the brain organization are unique.

### QPP4 and above

While we have described the three most prominent QPPs, the residuals after regression of these patterns can be reanalyzed to detect QPP4, and so on. Our additional analysis on forty randomly selected individuals showed QPPs 1-5 on average explain ~36% of the variance of the cortical parcels’ timeseries (QPPs 1-3 explain ~27%), with a wide range of ~8%-62% across parcels (~5%-52% for QPPs 1-3). QPP4 and above progressively explain less variance without a clear cutoff. Our focus on QPPs 1-3 was because they readily match the cortical FCGs 1-3 and account for a substantial portion of the functional connectivity.

### What is the underlying neuronal mechanism of the QPPs?

Multimodal studies in both animals (Pan et al., 2013; Thompson et al., 2014) and humans (Grooms et al., 2017) have shown that the QPPs detected with rsfMRI are linked to underlying infraslow electrical activity (loosely defined as < 1 Hz). Given the poorly characterized sources of infraslow activity, its relationship to the QPPs provides limited insight into the latter’s underlying mechanism. In fact, it may instead be that QPPs provide insight into the mechanisms underlying the infraslow electrical activity.

Coordinated activity throughout the whole brain, as demonstrated within QPPs, must occur over the framework of the brain’s anatomical architecture. One possibility then is that QPPs represent a sort of “resonance” of the brain’s structural network. This is inspired by a number of studies showing that the neural mass models, coupled with an estimated structural network obtained with diffusion-weighted MRI, can reproduce some aspects of the empirical rsfMRI-based functional connectivity (Deco et al., 2013; Cabral et al., 2017). However, when we applied QPP analysis to the output of these models (Kashyap and Keilholz, 2019), we found that while they capture some qualitative aspects of QPPs, such as the division of the brain into two anticorrelated clusters of areas, they are less successful at reproducing the complex dynamics observed within the QPPs, suggesting that other factors are involved.

Another neuronal mechanism proposed to underlie functional connectivity and therefore QPPs is spatially patterned input from subcortical neuromodulatory sources (e.g., the basal nucleus of Meynert (Chang et al., 2016; Liu et al., 2018b) or the rostral ventrolateral medulla (Drew et al., 2008)). These areas have widespread projections that differ in their density and/or receptor types, potentially allowing patterned modulation of ongoing brain activity that could manifest as QPPs. A growing body of work is exploring how neuromodulatory input affects functional connectivity. For example, Zerbi et al. (2019) recently showed that chemogenetic activation of the locus coeruleus (LC) increases the connectivity throughout the brain. Applying QPP analysis using the data and paradigm of such studies has the potential to reveal aspects about the neuronal mechanism underlying QPPs. Nevertheless, the old finding might turn out to be of particular importance. For example, Aston-Jones and Bloom (1981) has shown that the LC exhibits activity that varies in the infraslow range, matching the time scale of the QPPs. Moreover, since there are multiple QPPs, it is possible that different neuromodulatory sources dominate each QPP type, an intriguing avenue for future exploration.

### Timing and hemodynamics

Our results are based on timing differences in the blood oxygenation level dependent (BOLD) signals of different brain areas and regions. Because rsfMRI is sensitive to hemodynamics (instead of directly measuring neural activity), differences in neurovascular coupling, hence, hemodynamic response functions (HRFs) across areas could confound the timing differences attributed to their neuronal activity. If there is a gradual variation in the HRF’s time-to-peak across adjacent areas, the synchronous neural activity of these areas could result in a propagating BOLD signal across them. However, we do not think that the HRF’s difference is the dominant factor in the timing differences within QPPs. Within each QPP, the times of peak shift along functional gradients with consistent directions that match the known cortico-subcortical tract-based connections. Gradual/sharpness of such shifts between the same areas, and also the direction of such shifts, changes between QPPs. Blind deconvolution approaches have shown that variation in the HRF’s time-to-peak is around a second across the cortex (Wu et al., 2013), far shorter than the length of QPPs. Nevertheless, HRF variation could be a confound in our results, particularly in the propagation observed across some areas or the timing differences between regions. Future studies based on other more direct neuroimaging modalities, perhaps in rodents or other species with controlled settings, are necessary to validate our results and to address the open concerns.

### Possible task interactions

The RSNs that co-activate at different phases of the QPPs are closely related to networks activated by particular tasks (Smith et al., 2009). For this reason, it would be particularly interesting to examine the interaction between different tasks and QPPs. Some preliminary work shows that the spatial pattern of the QPP can be altered during task performance (Abbas et al., 2019a), but interpretation was limited by the use of existing data with short task blocks that makes it difficult to disentangle the effects of on and off blocks from the effects of task performance. It may also prove useful to minimize the effects of QPPs to isolate the activity that occurs in response to a task or stimulus. QPPs are such a prominent feature of rsfMRI data that they could easily obscure smaller changes related to localized neural activity. Regression of QPPs may also reduce the variability in response observed within and across individuals, an avenue worth future investigation.

### Clinical alterations and behavioral correlates

Functional connectivity is widely used as a biomarker in psychiatric disorders and neurological diseases. QPP analysis offers a tool to determine whether the alterations observed are localized to particular areas or connections, or more accurately depicted as downstream effects of disruption of the whole-brain pattern. For example, a number of neurological disorders exhibit disruption of DMN activity and early degeneration of brainstem nuclei like the locus coeruleus (Peterson and Li, 2018; Betts et al., 2019). In this case, the loss of functional connectivity in the DMN may be the result of a disruption of the brainstem and thalamic input that leads QPP1. If so, the strength, frequency, or spatial pattern of QPP1 may serve as more accurate biomarkers than measurements of the resulting functional connectivity. For example, we recently found that QPPs are weaker in ADHD patients than in healthy controls (Abbas et al., 2019b). Similarly, when looking for cognitive or psychological correlates, QPP-based measures may provide more information, or alternatively, QPPs may be minimized to emphasize remaining variability that may be more closely related to the relevant differences in neural activity. These questions have only begun to be explored.

## Conclusion

We have shown that functional connectivity predominantly arises from a few recurring spatiotemporal patterns of intrinsic activity, which sweep the macroscale FCGs in a coordinated way across the whole brain. These patterns can be obtained by simply averaging similar segments of rsfMRI timeseries and are robust to preprocessing choices. The neurophysiological mechanisms that underlie these patterns are still unknown. However, our results specifically suggest that thalamic and brainstem areas are key drivers for the intrinsic alternation of activity in the cortical nodes of the task positive and default mode networks. These patterns provide promising avenues for exploration in terms of clinical alterations and rest-task interactions. Taken with previous reports of QPPs in different species and under different conditions (Majeed et al., 2009; Belloy et al., 2018a, 2018b), our findings suggest that they reflect fundamental aspects of the brain’s functional organization.

## Acknowledgments

We thank Drs. Garth Thompson, Eric Schumacher, Anabelle Singer and Bruce Crosson for their very constructive comments as PhD committee members and all the previous and current lab members for the inspirational discussions. Since 2018, as we presented various parts of this work in multiple conferences, many researchers, including the authors of our key references, gave influential comments and we sincerely thank them all. We would also like to acknowledge our funding sources for this work, NIH R01MH111416 and R01NS078095.

