## Supplementary material for "Propagating patterns of intrinsic activity along macroscale gradients coordinate functional connections across the whole brain": S.M.

**Supplemental Material: Statistical Evaluation of Activity within QPPs**

*1.1. Definition of significant activity within a QPP*. To quantitatively analyze the activity within a QPP, only the vertices/voxels that exhibit statistically significant activation/deactivation, at any time interval within the ~20s duration of that QPP (for brevity, the active vertices/voxels) were included. To identify these active vertices/voxels, first, we built null spatiotemporal patterns by averaging randomly selected Ni number of non-overlapping segments, from all four scans of all 817 individuals, with Ni being the number of the contributing segments to the group QPPi (i=1:3). The random selection was repeated 50 times for each Ni, resulting in 50 null patterns. Next, we calculated the root sum square of each ~20s-long timecourse corresponding to each vertex/voxel within each null pattern and found the $99^{th}$ percentile of the root sum square values across all vertices/voxels and all null patterns (~92K×50 entries; Fig.S7). This percentile, which can serve as a power threshold to determine the active vertices/voxels of a QPP, increases as the number of randomly selected segments decreases, i.e., it is higher for N3 segments (corresponding to QPP3) compared to N2 or N1 because there are fewer segments that contribute to QPP3. To simplify, we used the threshold obtained based on N3 randomly selected segments for all QPPs. Finally, we identified the active vertices/voxels of each QPP as those vertices/voxels whose timecourse has a root sum square larger than the threshold.

Note, usage of root sum square instead of root mean square (rms) to indicate the power of a timecourse, was for the slight improvement in the computational precision given that all the timecourses in the null patterns and QPPs have equal number of timepoints (i.e., 30 timepoints). Furthermore, the following is the rational for defining significant activity as described above. If a QPP is an average of certain similar segments, then any of its timecourses with a power near that of a null pattern, built by averaging randomly selected segments (that are not necessarily similar and therefore should average out to near zero), is basically due to properties of rsfMRI timeseries alone. If the power of a QPP's timecourse is well larger than that of a null pattern, e.g., above $99^{th}$ percentile, then we can say that timecourse entails significant activity.

*1.2. Timepoints of significant activity within a QPP*. To determine the timepoints that the active vertices/voxels exhibit statistically significant activation/deactivation within the 30-timepoint duration of a QPP, we found the $99^{th}$ percentile of the magnitudes (i.e., the absolute values) of all 30 timepoints per timecourse of all vertices/voxels and all null patterns corresponding to each Ni

(~92K×30×50 entries for each Ni; Fig.S8). This percentile also increases as Ni decreases and to simplify, we again used the N3-based threshold to serve as the magnitude threshold for all QPPs.

We further added a second step for the identification of the active vertices/voxels of a QPP by only including the vertices/voxels with peak magnitude (i.e., either the magnitude of peak activation or the magnitude of dip deactivation) larger than the magnitude threshold. Finally, timepoints of significant activity of active vertices/voxels can be find as those timepoints at which the magnitude of the timecourse is larger than the magnitude threshold.

*1.3. Summary maps of activity within a QPP: primary basis for further statistical analysis.* Only including the active vertices/voxels, the summary maps of the activity within each QPP were calculated, by clustering the timecourses into ten clusters and ordering them (coarse summary), described in detail in the main text, and finding the time of peak activation and the time of dip deactivation for each timecourse (fine summary). These summary maps are the primary basis for further statistical analysis. For the fine summary map, we only included the times that correspond to a peak/dip larger than the magnitude threshold described previously. We included the time of dip because of a few clusters of timecourses with only supra-threshold dip but not supra-threshold peak due to their particular phase. If the QPP is phase-adjusted to have a reversed phase, the same timecourses exhibit supra-threshold peak but not supra-threshold dip.

*2. Significant coordination of activity within a QPP*. To quantitatively show the coordinated propagation of activity across the whole brain within a QPP, we separated the ten clusters of the QPP's timecourses into seven brain regions (cerebral cortex, cerebellum, thalamus, hippocampus, amygdala, brainstem and deep brain nuclei, striatum), resulting in seventy groups, and tested the significance of differences in the times of peak activation and the times of dip deactivation between pairs of groups, using t-test (both dependent and independent) and corrected alpha value of 0.01/(70×69/2)=4.1e-6. Since some of the groups have a few or even no members, we excluded the groups whose size are lower than 15 ($5^{th}$ percentile across all 70 groups). To address the difference in size for any pair of groups (cortical regions are much larger than non-cortical regions), we lowered the size of the larger group by random selection, repeating and testing 50 times and choosing the maximum p-value across the repetitions. As supplementary tests, we also performed independent t-test and Kolmogorov-Smirnov test (ks-test), without matching the group sizes, and both tests had similar outcome compared to using the dependent t-test.

The results using only dependent t-test are brought in three supplementary figures for QPPs 1-3 (Fig. S12, S18, S21, respectively); each of these figures shows the timecourses per each included group (out of 70), along with the size and median of time-of-peak for that group, and finally, the matrix of significant difference in time-of-peak between pairs of the included groups. Each supplementary figure along with our main figure which shows the maps of a QPP's clusters (Fig.2) are the statistical support for our statements about the propagation of activity in each brain region; specifically, the order of the clusters of timecourses that spatially tile a region along with the significant progression of their median time-of-peak support the propagation of activity.

The coordinated activity observed within a QPP also involves a time-locked activity between two non-adjacent areas, e.g., a cortical and a non-cortical area. Such time-locked activity, or coactivity, is supported by the two areas belonging to the same cluster of timecourses, despite possible slight but significant time differences. The supplementary figures described above (Fig.S12, S18, S21) along with the maps of clusters (Fig.2) also support coactivity between non-cortical and cortical areas stated in our report (summarized in Table 1).

There are instances of focal propagation of activity specifically mentioned for QPP1 across V1 and across the lateral temporal lobe (TL). The sweep across TL seems to be time-locked with the posterior-anterior sweep along the hippocampus towards the amygdala because of the same clusters of timecourses with the same order that tile these areas. To quantitatively show the occurrence of these focal sweeps and the abovementioned coactivity, we tested the significance of time differences between the clusters in the V1 and also the between clusters in the TL and the hippocampus-Amygdala (considered as one region, abbreviated as H&A), using the same procedure described in the first paragraph of the current part. For V1, we only tested the time of peak activation and for the TL and H&A, we only tested the time of dip deactivation, but we separated the left and right hemispheres. The results are brought in a supplementary figure (Fig.S15) with a similar format described in the previous two paragraphs.

*3.1. Comparison between QPPs and FCGs*. To quantitatively show that the propagation of activity within a QPP is also along a particular functional connectivity gradient (FCG) across the cortical sheet, we calculated the correlation between the QPP's time of peak map and that FCG. We obtained the cortical FCGs from a dscalar.nii file published in Margulies et al (2016) and downloadable from the first author's website. The cortical vertices that were identified as non-active within the QPP or those with the peak magnitude smaller than the threshold, which overall are only ~2% of ~60K cortical vertices, were excluded from the FCG vector before correlating it with the QPP's time of peak map. We also found the correlation map within the QPP, with Left PCC (or left V2 for QPP3) being the seed timecourse, and correlated this correlation map with its corresponding FCG, which resulted in slightly higher correlation compared to QPP's time of peak map.

The results are brought in three supplementary figures for QPPs 1-3 (Fig. S13, S19, S22, respectively, parts a-b), where maps of time of peak and correlation for each QPP are plotted aside a particular cortical FCG and the correlation values are brought in the caption.

Although we only qualitatively compared each QPP with the existing non-cortical FCGs for the three regions of the cerebellum, hippocampus and striatum, the statistically evaluated summary map of each QPP per each region was the basis for such comparison.

*3.2. Timing differences across cortical RSNs within QPPs*. To quantitatively show the sequential activity of RSNs within a QPP (which is the basis for our description of QPP's activity and also another way of showing the propagation of activity within a QPP), we grouped the active cortical vertices of that QPP according to the Yeo's seven cortical RSNs and tested the significance of differences in the time of peak activation between pairs of groups, using t-test (both dependent and independent) and corrected alpha value of $0.01/(7\times6/2)=4.8e\text{-}4$. To address the group size difference between any pairs of groups, we reduced the size of the larger group to the size of the smaller group by random selection, repeating such random selection plus the t-test 50 times and choosing the maximum p-value across the repetitions (independent t-test and ks-test, without matching group sizes, were performed and resulted in similar outcome).

Moreover, we averaged the QPP's timecourses across each group (i.e., RSN), resulting in seven timecourses corresponding to the seven cortical RSNs and calculated the $7\times7$ correlation matrix between all pairs of timecourses to support a few statements.

Although only the active cortical vertices were included, we also calculated a magnitude threshold for activation/deactivation of the average timecourse across a cortical RSN. We averaged the timecourses of each null pattern (described in part 1.1) across the cortical RSNs and found the 99$^{th}$ percentile of magnitude values of all seven timecourses across all 50 null patterns (Fig.S14a). In all QPPs, timecourses of all seven cortical RSNs reach above the magnitude threshold, hence, entail significant activity.

Finally, as a supplementary analysis, we also found the time of zero-crossing, from active to deactive or vice versa, for the active cortical vertices belonging to the seven RSN groups and tested the significance of differences between pairs of groups, using the same procedure described above for the time of peak. This analysis was to test the observation that a few RSNs seem to remain active/deactive as the sign of activity is switching in other RSNs.

The results of the current part, using only dependent t-test, are brought in three supplementary figures for QPPs 1-3 (Fig. S13, S19, S22, respectively, parts c-e); each of these figures shows seven timecourses, each timecourse being the average of the QPP's timecourses across a cortical RSN (i.e., group), along with the 7×7 correlation matrix between pairs of timecourses, the median times of peak and zero-crossing per group and matrices of significant difference between pairs of groups in terms of times of peak and zero-crossing. Significant progression of times of peak and zero-crossing of QPP1's timecourses when grouped according to cortical RSNs supports the sequential activity of cortical RSNs within QPP1 (summarized in Table 1). Note, based on the consistencies of propagation (Table 2), we first came up with a summary (Table 1), then we performed the analysis of this part, which supports that summary.

*3.3. QPPs versus the existing non-cortical parcellation schemes.* To compare QPP's coarse summary of activity with the existing parcellation schemes (Table S1), particularly at the non-cortical regions, we found the size of each cluster of QPP's timecourses per network/parcel/ probabilistic-map per region. Results are brought in three supplementary tables for QPPs 1-3 (Table S2-4, respectively). For the cerebellum and striatum (and also for the cerebral cortex), such quantification supports the overall match between the activity within QPP1 and the adopted functional networks. This match is mostly based on the first two clusters of QPPs, which are the largest and are anticorrelated with one another. For other non-cortical regions, the provided sizes mostly serve to identify which anatomical areas contain which clusters.

We did not group the non-cortical voxels according to the parcellation schemes to test for the timing differences, similar to what we have done for the cortical RSNs. The reason is the overall less consensus about the existing non-cortical parcellation schemes and the observation that the coactivity map of non-cortical areas with the cortical areas changes between QPPs (for example, in two regions of the brainstem and deep brain or the striatum, compare QPP1 and QPP2, using the cluster maps in Fig.2b and 2e or Tables S2 and S3; also note our discussion under the first subheading, "functional connectivity, RSNs and FCGs", in the fourth paragraph) - parcellation methods based on static functional connectivity could easily not be sensitive enough to such dynamic changes, reducing the consistency with subtle changes in method.

*4. Timing differences between brain regions within QPPs.* To examine the nuanced timing differences between brain regions during a QPP, that can suggest the driving mechanisms, per each of seven regions, we found the number of vertices/voxels with a significant peak at each timepoint of a QPP, resulting in seven histograms, each with 30 bins corresponding to the QPP's timepoints (brought in Fig.3). Since QPPs 1-3 all involve two large anticorrelated clusters of areas, their distributions of time of peak per brain region (or across the whole) are bimodal. For the scope of this work, we only focused on the second mode, identified as the entries above the mid timepoint of the cortical distribution. Therefore, first, we found the mid timepoint of the cortical distribution. Then, for each of seven brain regions, we found all the time of peaks higher than that mid timepoint and tested the significance of differences between regions using the same procedure described earlier (using dependent t-test and matching group sizes; independent t-test and ks-test, without matching group sizes, had similar outcome). Only the time of supra-threshold peak of the active vertices/voxels were included and alpha value was corrected for multiple comparisons ($0.01/(7\times6/2)$). Results are brought in a supplementary figure (Fig.S17) that shows median of time of peak per region and the significance matrix between pairs of regions for each QPP.
