## Supplementary material for "Propagating patterns of intrinsic activity along macroscale gradients coordinate functional connections across the whole brain": Fig.S

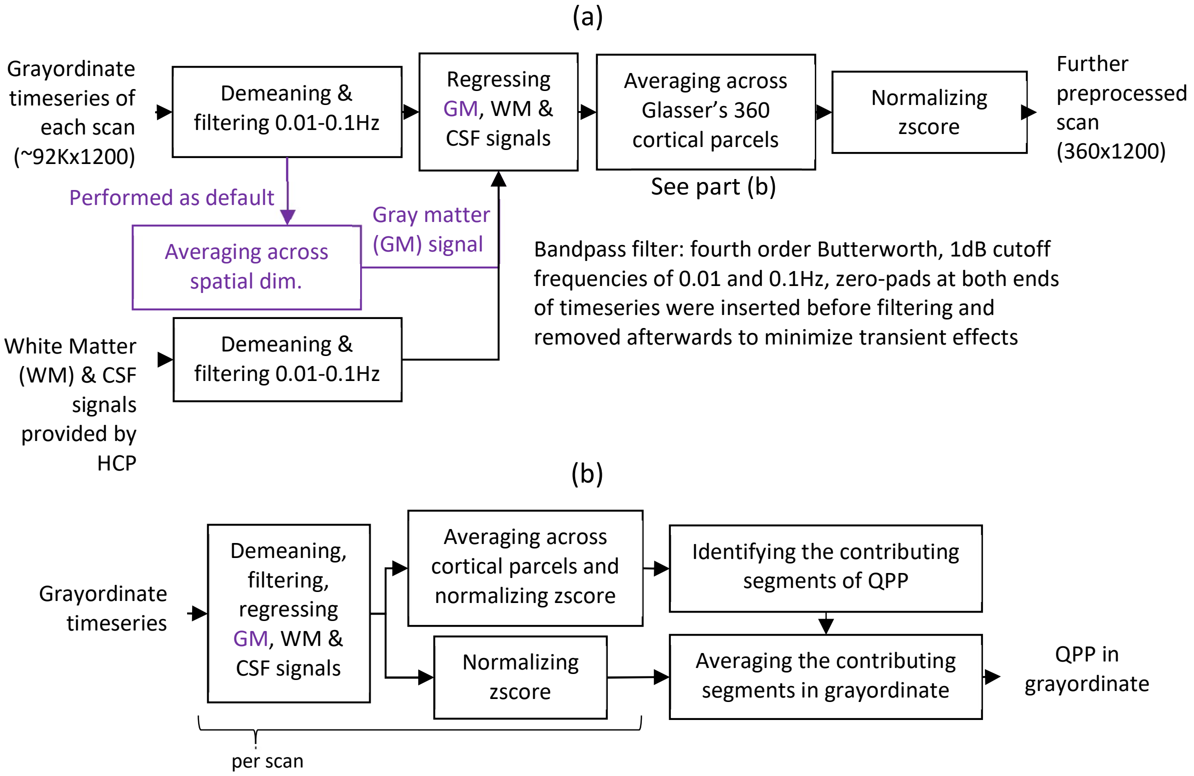


Fig.S1 (a) Additional preprocessing applied to the minimally preprocessed grayordinate and FIX denoised rsfMRI scans of HCP S900 dataset. (b) To compensate the parcellation used in (a), we can obtain a QPP in grayordinate when its contributing segments are identified based on a rsfMRI scan in parcel-space.

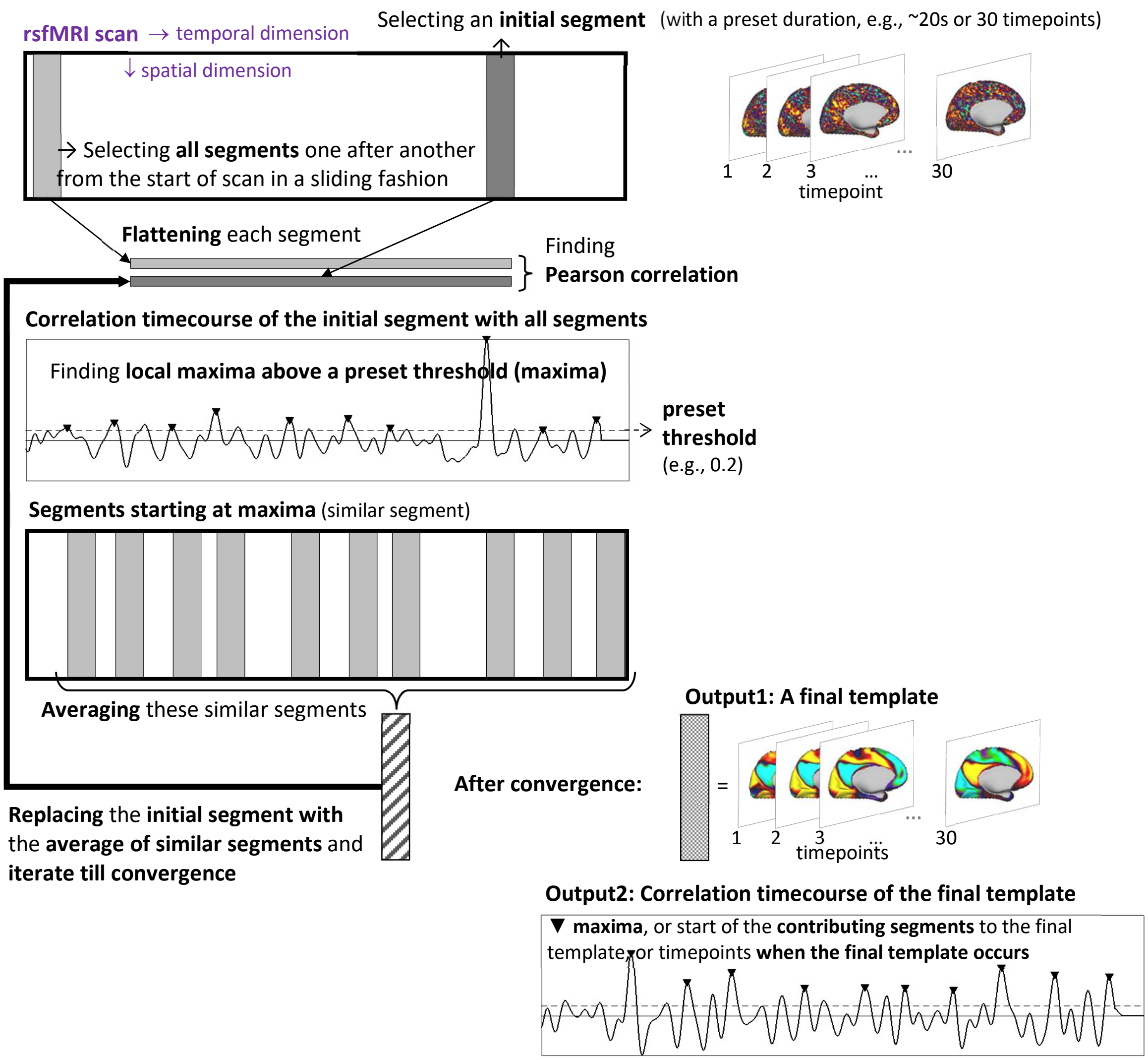


Fig.S2 The main algorithm to detect a QPP, developed by Majeed et al., 2011, is correlation-based and iterative. It identifies similar segments of a functional scan and averages them for a representative spatiotemporal template.

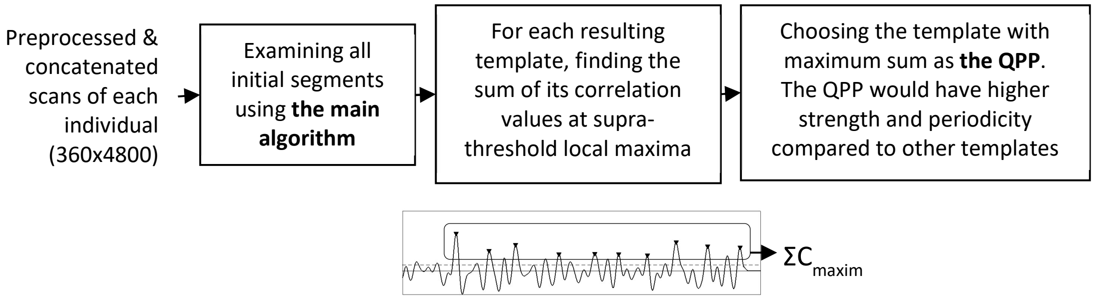


Fig.S3 Robust detection of the QPP for an individual, introduced in Yousefi et al., 2018.

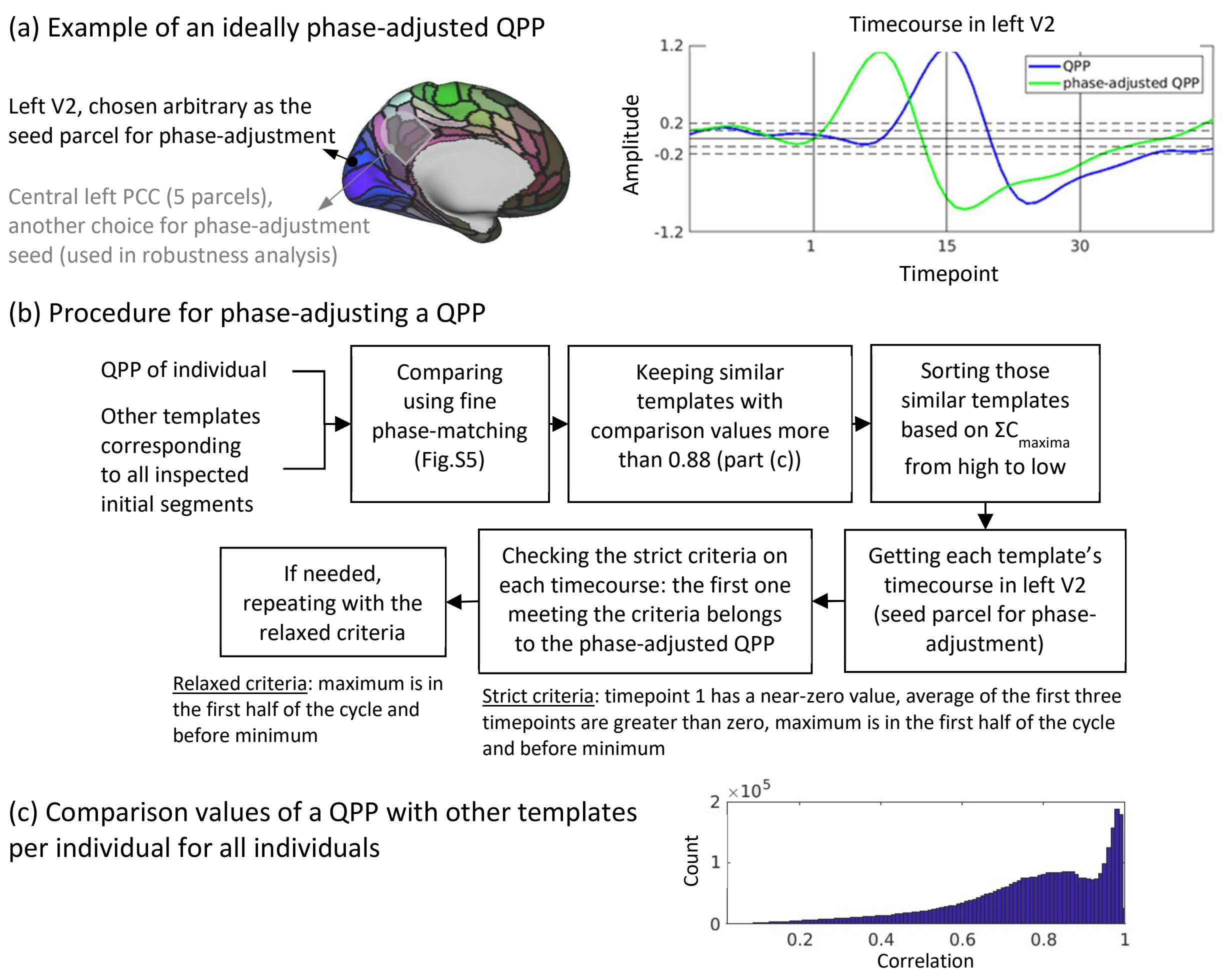


Fig.S4 (a) Example of an ideally phase-adjusted QPP. (b) Procedure for phase-adjusting a QPP. (c) Comparing QPP of an individual with other templates results in a bimodal distribution, when combining all individuals, with lower mode (0.88) taken as the threshold in part (b).

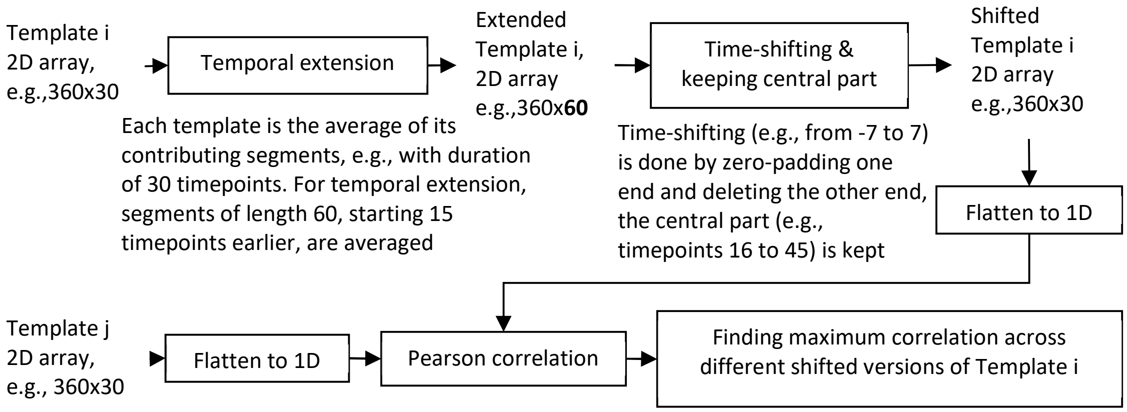


Fig.S5 Comparing two templates by fine phase-adjusting (Template i = QPP1 for phase-adjusting).

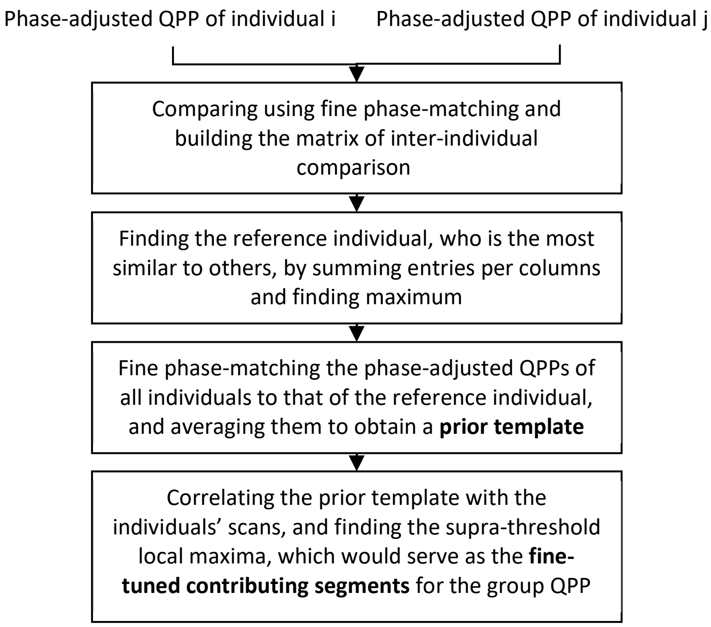


Fig.S6 Obtaining the group QPP by fine-tuned averaging.

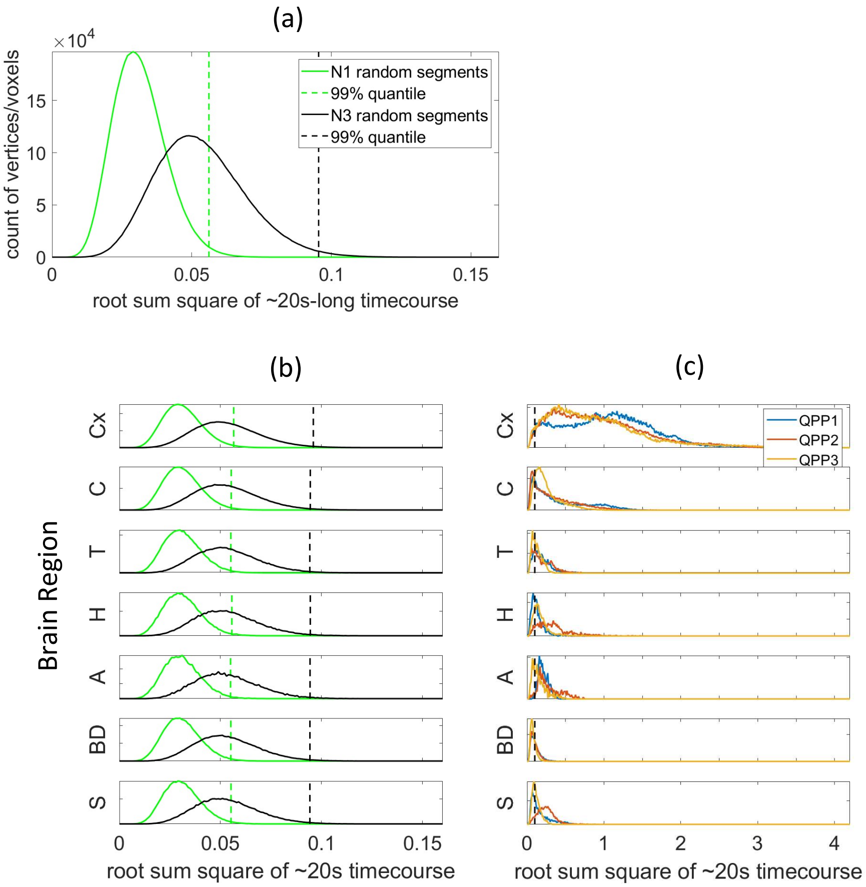


Cx: Cerebral cortex, C: Cerebellum, T: Thalamus, H: Hippocampus, A: Amygdala, BD*: Brainstem and Deep brain, S*: Striatum
*As anatomically labeled in the HCP dataset in grayordinates, we have included Brainstem, Ventral Diencephalon and Pallidum in BD, and Caudate, Putamen and Accumbens in S.

Fig.S7 Statistics to identify the active vertices/voxels of a QPP (i.e., vertices/voxels that become activated/deactivated at any time interval within the ~20s duration of that QPP; corresponding to Supplementary Material, S.M., part 1.1). (a) Histograms of root sum square of ~20s-long timecourse corresponding to a vertex/voxel within a null pattern, across all ~92K vertices/voxels and all 50 null patterns (~92Kx50 entries; note, root sum square of a timecourse is used to indicate the power of that timecourse instead of root mean square, given that all timecourses have equal length); each null pattern was built by averaging randomly selected N1 (~28.3K) or N3 (~9.8K) number of segments, with Ni being the number of the contributing segments to the group QPPi. 99th percentile of root sum square values, which can serve as a power threshold to identify active vertices/voxels, increases as Ni decreases; to simplify, we used the threshold obtained based on N3 randomly selected segments (0.1) for QPPs 1-3. (b) Histograms shown in part (a) separated for each brain region, showing near identical distributions for all regions. (c) Histograms of root sum square of timecourses for QPPs 1-3 per brain region vs the threshold for identifying the active vertices/voxels (0.1); as expected, the root sum square values in the subcortical regions are lower compared to the cerebral cortex.

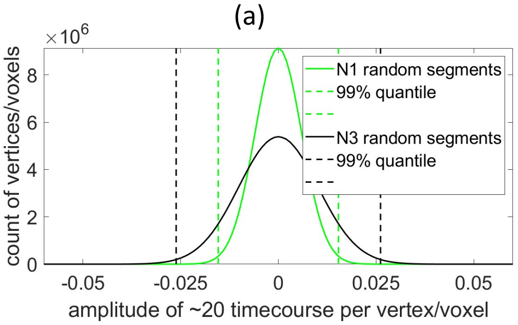


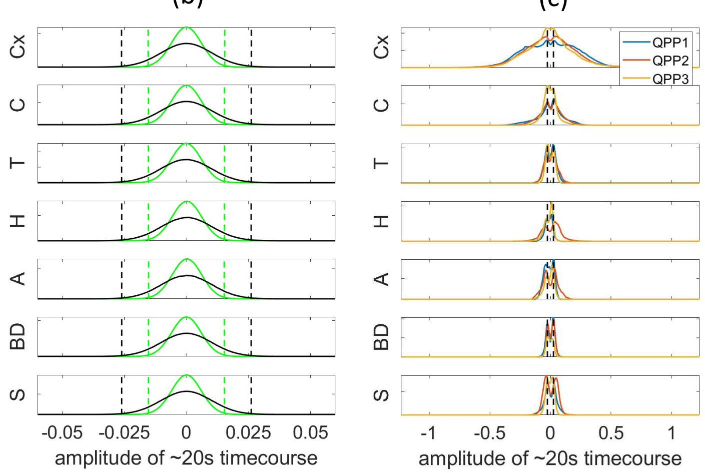


(d)

| | | QPP1 | QPP2 | QPP3 | All |
|---|---|---|---|---|---|
| **Number of active vertices/voxels** | **Cx** | 58317 | 58452 | 58088 | 59412 |
| | **C** | 14526 | 14248 | 15672 | 17853 |
| | **T** | 1939 | 1836 | 1500 | 2536 |
| | **H** | 912 | 1439 | 1215 | 1559 |
| | **A** | 639 | 617 | 422 | 647 |
| | **BD** | 2349 | 2252 | 1287 | 5447 |
| | **S** | 2349 | 3535 | 2195 | 3828 |
| | **All** | 81031 | 82379 | 80379 | 91282 |
| **Percent of active vertices/voxels** | **Cx** | 98 | 98 | 98 | |
| | **C** | 81 | 80 | 88 | |
| | **T** | 76 | 72 | 59 | |
| | **H** | 58 | 92 | 78 | |
| | **A** | 99 | 95 | 65 | |
| | **BD** | 43 | 41 | 24 | |
| | **S** | 61 | 92 | 57 | |
| | **All** | 89 | 90 | 88 | |

Cx: Cerebral cortex, C: Cerebellum, T: Thalamus, H: Hippocampus, A: Amygdala, BD: Brainstem and Deep brain, S: Striatum

Fig.S8 Statistics to determine the activation/deactivation time intervals within the ~20s duration of a QPP (corresponding to S.M.1.2). (a) Histograms of the amplitude value at each timepoint of each timecourse corresponding to a vertex/voxel within a null pattern, across all timepoints, timecourses and null patterns (30x~92Kx50 entries); each null pattern was built by averaging randomly selected N1 (~28.3K) or N3 (~9.8K) number of segments, with Ni being the number of the contributing segments to the group QPPi. 99$^{th}$ percentile of the magnitude values (i.e. the absolute of the amplitude values), which can serve as a threshold to define activation/deactivation time intervals, increases as Ni decreases; to simplify, we used the threshold obtained based on N3 randomly selected segments (0.025) for QPPs 1-3. (b) Histograms shown in part (a) separated for each brain region showing near identical distributions. (c) Histograms of amplitude value of only the active vertices/voxels for QPPs 1-3 vs the magnitude threshold; amplitude values in the subcortical regions are lower compared to the cerebral cortex, as expected. A second step for the identification of the active vertices/voxels of a QPP was added by only including the vertices/voxels with peak magnitude (i.e., amplitude of peak activation or dip deactivation) larger than the magnitude threshold. (d) Total number of active vertices/voxels and their percentage per region.

Table S1 Existing parcellations, functional networks and gradients* for comparison with QPP

| Region | Parcellations and gradients | References |
|---|---|---|
| Cortex | Seven resting-state networks (RSNs), the first three functional connectivity gradients (FCGs) and 360 multimodal parcellation to identify cortical areas | Thomas Yeo et al., 2011; Margulies et al., 2016; Glasser et al., 2016 |
| Cerebellum | RSN-based parcellation and two cerebellar FCGs | Buckner et al., 2011; Guell et al., 2018 |
| Thalamus | Tractography-based parcellation | Behrens et al., 2003 |
| Hippocampus | Multimodal parcellation and two hippocampal FCGs | Robinson et al., 2016; Vos De Wael et al., 2018 |
| Amygdala | Structural parcellation** | Tyszka and Pauli, 2016 |
| Brainstem and deep brain nuclei+ | Probabilistic maps of locus coeruleus (LC), ventral tegmental area (VTA), substantia nigra pars compacta/pars reticulata (SNc/SNr), basal nucleus of Meynert (BNM), pedunculopontine nucleus (PPN), dorsal raphe (DR), median raphe (MR), periaqueductal gray (PAG) ***, subthalamic nucleus (STH), globus pallidus external/internal (GPe/GPi) and hypothalamus (HTH) | Keren et al., 2009 (LC); Pauli et al., 2016 (VTA, SN, GP, STH, HTH); Li et al., 2014 (BNM); Edlow et al., 2012 (PPN, DR/MR, PAG) |
| Striatum+ | RSN-based parcellation and two striatal FCGs | Choi et al., 2012; Marquand et al., 2017 |

* Other than the cortical gradients, we did not obtain any file for the non-cortical regions and just qualitatively compared the non-cortical gradients with the statistically evaluated summary maps of QPPs.

** After downloading the amygdala parcellation, to simplify, we regrouped the parcels into six (P1 to P6), which are as follows. P1: lateral (La), P2: basolateral (BL), P3: accessory basal (BM), P4: Paralaminar (BLV), P5: corticomedial nucleus (CMN) and central nucleus (CEN), P6: amygdalostriatal transition area (ASTA), periamygdaloid cortex (ATA), anterior amygdaloid area (AAA) and intercalated nuclei (AMY).

*** DR, MR and PAG are medial nuclei, not showing up in the representative planes chosen for the main figures.

\+ As anatomically labeled in the HCP grayordinates, we included the Brainstem, Ventral Diencephalon and Pallidum in the "Brainstem and deep brain region" and the Caudate, Putamen and Accumbens in the "Striatum region".

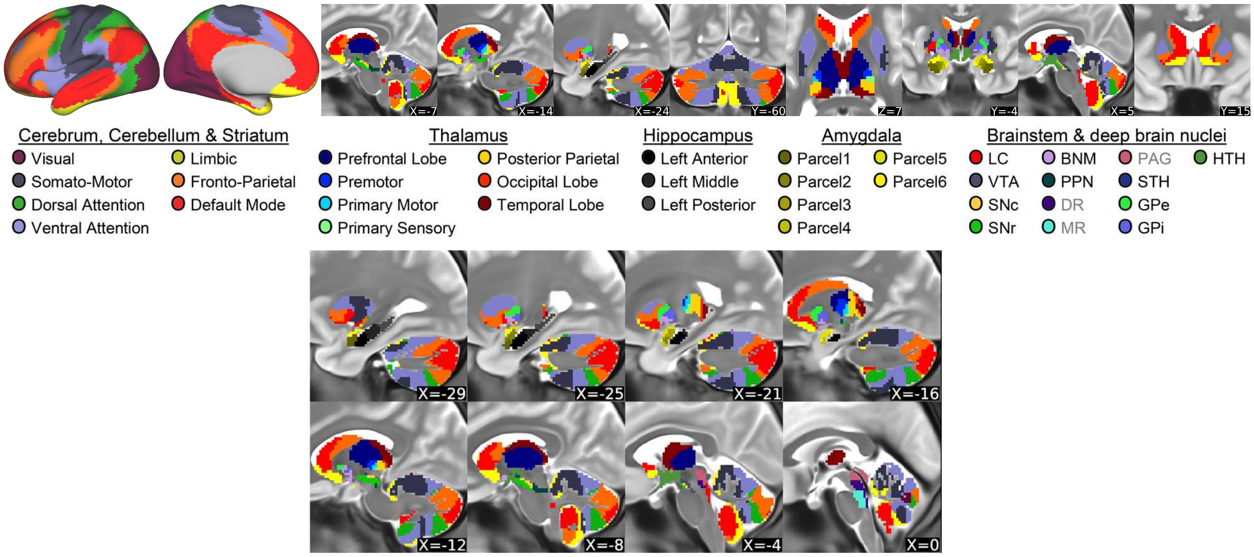


Fig.S9 Existing parcellations and functional networks. Representative planes chosen for the main figures (top part) and more sagittal planes (bottom part)

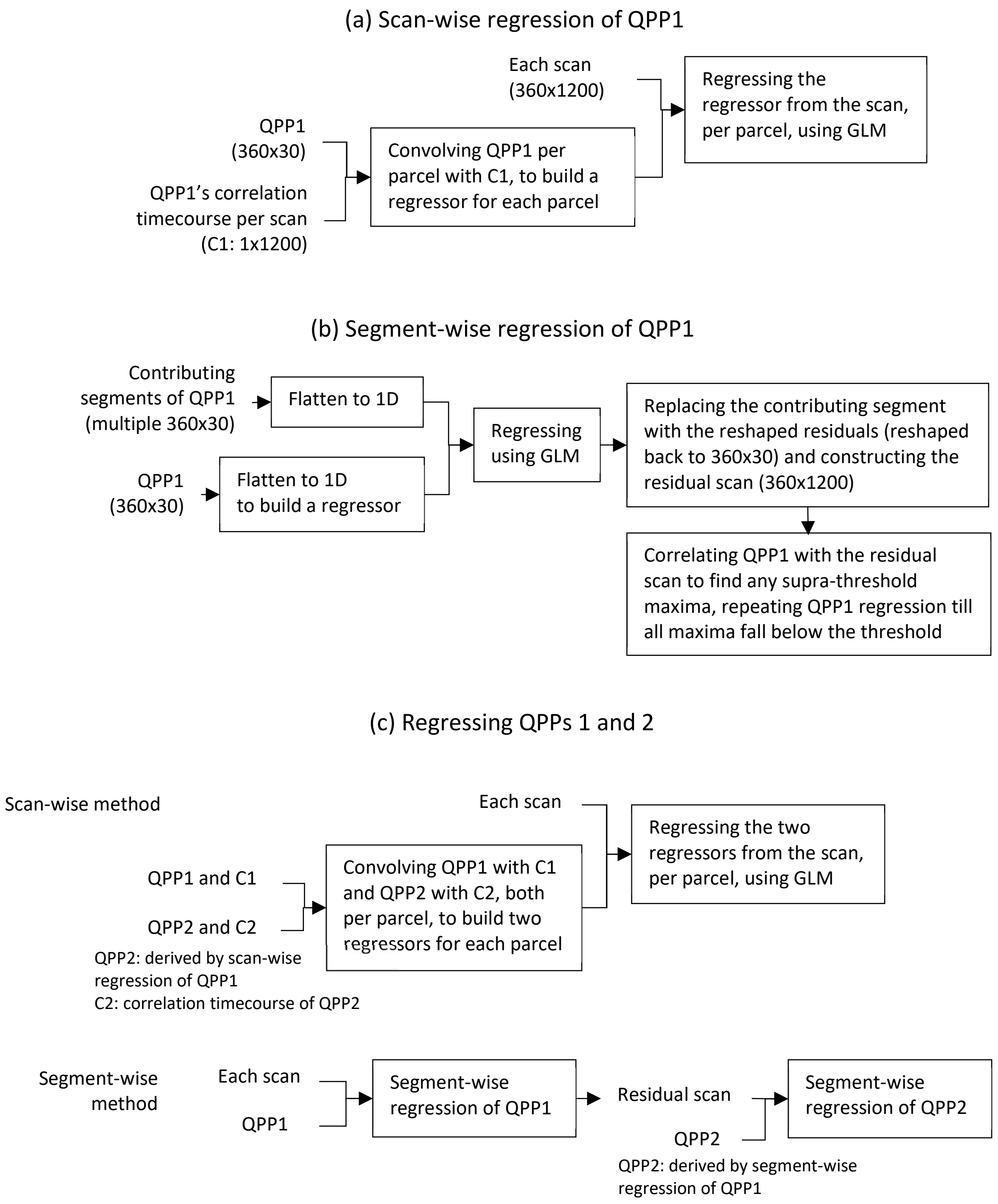


Fig.S10 Scan-wise (a) and segment-wise (a) regression of QPP1. Regressing QPPs 1 and 2 (c).

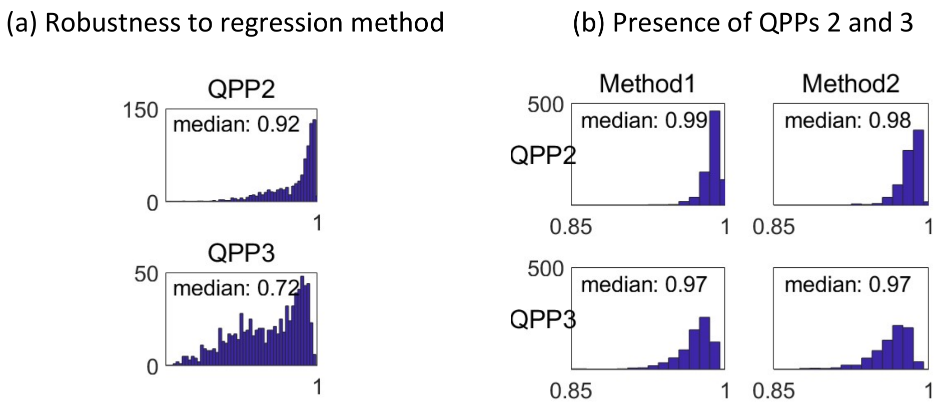


Fig.S11 (a) Correlation between QPPs 2 and 3 of individuals obtained by two regression methods. (b) Correlation between QPPs 2 and 3 of individuals obtained based on the original scans and residual scans using both regression methods. Nearly identical results show QPPs 2 and 3 are present in rsfMRI timeseries, similar to QPP1.

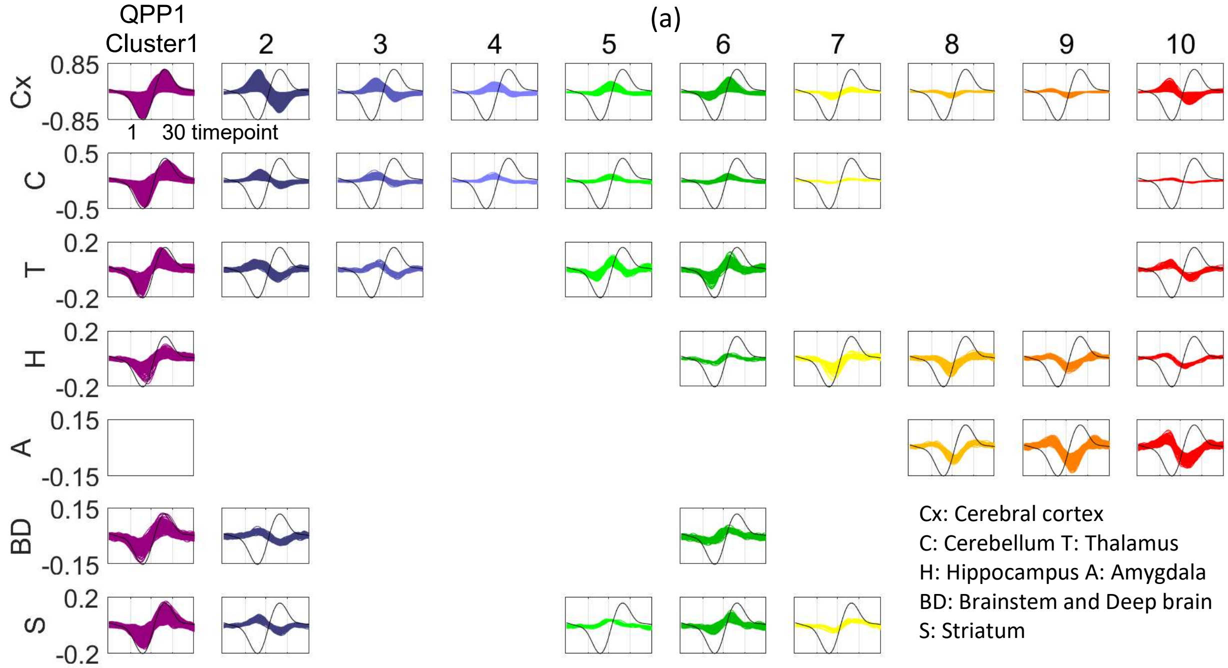


(b) Median time of peak

| | Cx | C | T | H | A | BD | S |
|---|---|---|---|---|---|---|---|
| **1** | 24 | 25 | 21 | 24 | - | 23 | 25 |
| **2** | 9 | 11 | 9 | - | - | 10 | 10 |
| **3** | 13 | 13 | 13 | - | - | - | - |
| **4** | 16 | 15 | - | - | - | - | - |
| **5** | 17 | 17 | 18 | - | - | - | 17 |
| **6** | 19 | 19 | 19 | - | - | 19 | 20 |
| **7** | 25 | - | - | 25 | - | - | 25 |
| **8** | 25 | - | - | - | - | - | - |
| **9** | 4 | - | - | 4 | 5 | - | - |
| **10** | 7 | - | 8 | 4 | 5 | - | - |

(c) Significance of difference in time of peak
significant non-significant (p=4e-6)

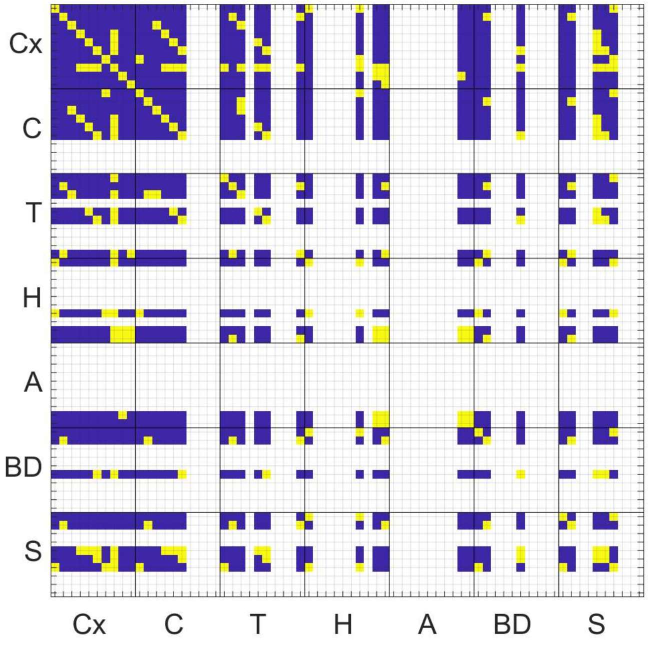


(d) Cluster size

| | Cx | C | T | H | A | BD | S |
|---|---|---|---|---|---|---|---|
| **1** | 18935 | 8795 | 750 | 267 | - | 1909 | 1755 |
| **2** | 30572 | 3115 | 389 | - | - | 180 | 125 |
| **3** | 1757 | 1340 | 50 | - | - | - | - |
| **4** | 579 | 392 | - | - | - | - | - |
| **5** | 613 | 306 | 132 | - | - | - | 28 |
| **6** | 1260 | 383 | 431 | 17 | - | 148 | 118 |
| **7** | 876 | 24 | - | 121 | - | - | 32 |
| **8** | 397 | - | - | 250 | 49 | - | - |
| **9** | 319 | - | - | 155 | 341 | - | - |
| **10** | 2325 | 17 | 53 | 22 | 235 | - | - |

Fig.S12 Support for the propagation of activity within QPP1, without relying on FCGs and RSNs, and also support for the coactivity of non-cortical areas with cortical areas within QPP1 (both correspond to S.M.2). (a) QPP1's clusters of timecourses per brain region (Fig.2b shows the map of these clusters; LPCC timecourse plotted in black for reference). (b) Median time of peak per cluster per region. (c) Significance of differences in the time of peak between included clusters in all regions, (continued)

(e) Median time of dip

| | Cx | C | T | H | A | BD | S |
|---|---|---|---|---|---|---|---|
| **1** | 9 | 10 | 8 | 9.5 | - | 8 | 9 |
| **2** | 23 | 26 | 24 | - | - | 25 | 24 |
| **3** | 27 | 27 | 27 | - | - | - | - |
| **4** | 28 | 29 | - | - | - | - | - |
| **5** | 4 | - | 5 | - | - | - | - |
| **6** | 6 | 5 | 7 | - | - | 5 | 8 |
| **7** | 13 | 12 | - | 12 | - | - | 13 |
| **8** | 15 | - | - | 15 | 16 | - | - |
| **9** | 16 | - | - | 17 | 18 | - | - |
| **10** | 20 | - | 21.5 | 18 | 19 | - | - |

(f) Significance of difference in time of dip
significant non-significant (p=4e-6)

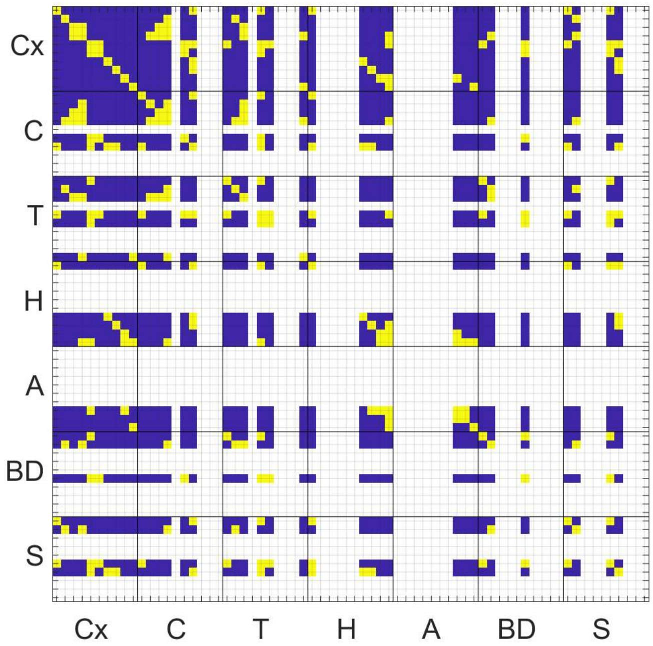


using dependent t-test (p=0.01/(70x69/2)); to match cluster size (shown in (d)) for the t-test, the size of the larger cluster was reduced by random selection 50 times, and the highest p-value was taken (included clusters had above 15 members). The propagation of activity per region within QPP1 is supported by the significant progression of the median time-of-peak across clusters of QPP1's timecourses when taken together with the spatial order of these clusters (Fig.2b). Furthermore, the coactivity of non-cortical areas with cortical areas, as stated in our report (summarized in Table 1) is supported by those noncortical and cortical areas belonging to the same cluster, despite possible slight but significant time differences (for example, the cerebellar or thalamic areas of cluster 1 have slight but significant time differences with the cortical areas of cluster 1, still they can be called coactive with those cortical areas since they all belong to the same cluster) . (e) Median time of dip of clusters (shown in part (a)) and (f) Significance of differences in the time of dips (obtained similar to part (c)).

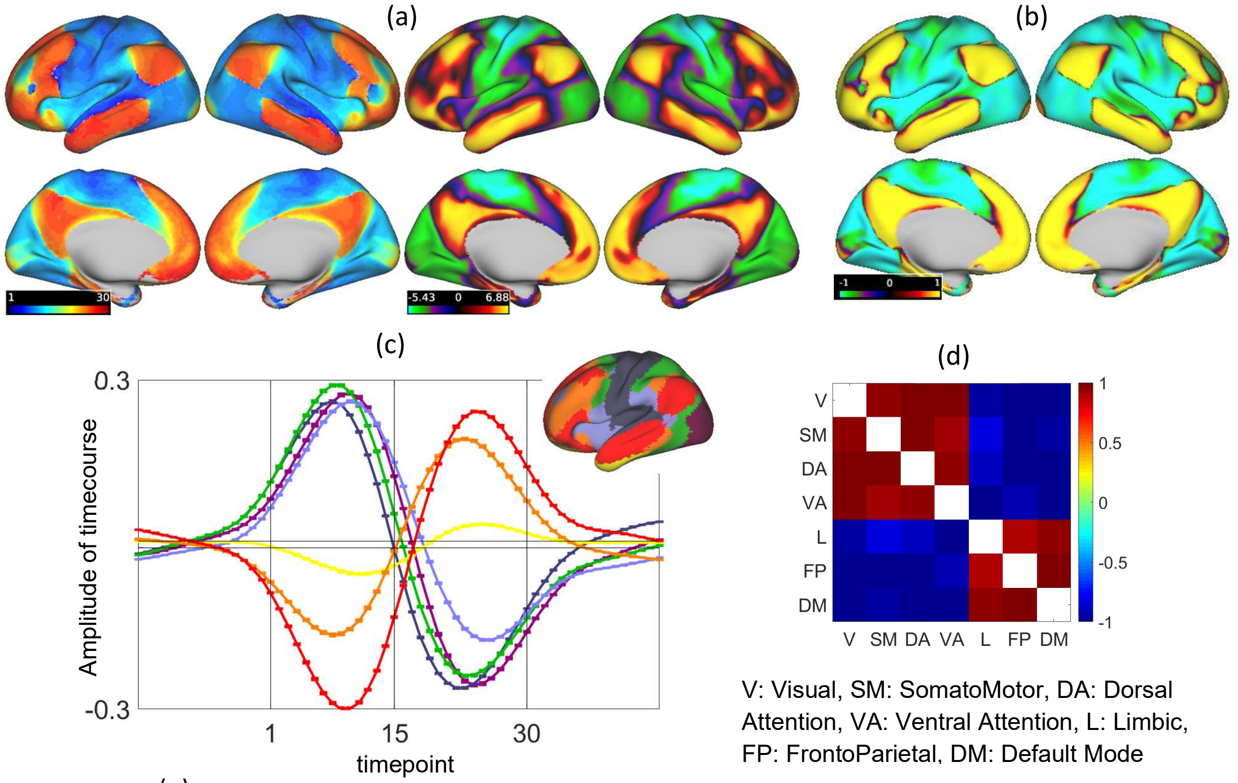


(e)

| | Time of peak | Time of zero-crossing | Number of active vertices |
|---|---|---|---|
| V | 10 | 16 | 8715 |
| SM | 8 | 14 | 11960 |
| DA | 8 | 15 | 6730 |
| VA | 10 | 18 | 7151 |
| L | 24 | 17 | 3785 |
| FP | 23 | 16 | 7254 |
| DM | 24 | 17 | 12722 |

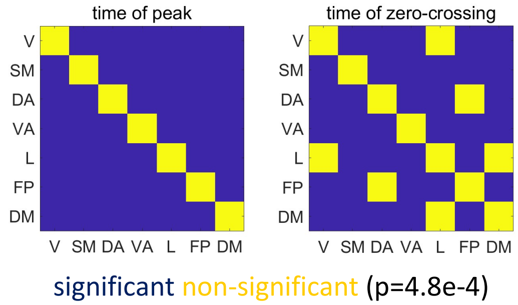


Fig.S13 Supports for the propagation of activity along cortical FCG1 and sequential activity of cortical RSNs within QPP1 (corresponding to S.M.3.1 and 3.2). (a) QPP1's time of peak map versus FCG1. The correlation between these two maps is 0.91, supporting that the peak activity within QPP1 sweeps FCG1. (b) Correlation map within QPP1, with LPCC being the seed timecourse, is also similar to FCG1 (correlation: 0.92). (c) Seven timecourses, each the average of the QPP1's timecourses across a cortical RSN (map of RSNs in the top right serves as the legend; all timecourses entail significant activity, see Fig.S14a; error bars show the standard error). (d) Correlation between timecourses shown in (c). (e) Median of time of peak and time of zero-crossing per group (i.e., RSN). Significant progression of times of peak and zero-crossing of QPP1's timecourses when grouped according to cortical RSNs supports the sequential activity of cortical RSNs within QPP1, which can be summarized as SM→DA & VA→FP/DM (Table1). For significant testing here, we used dependent t-test (p=0.01/(7x6/2)=4.8e-4) and for matching the group size for each pair of groups, we randomly sampled the larger group, repeated 50 times and took the highest p-value across repetitions. As an interesting note, VA is the last network to switch, i.e., remains active as other RSNs are switching.

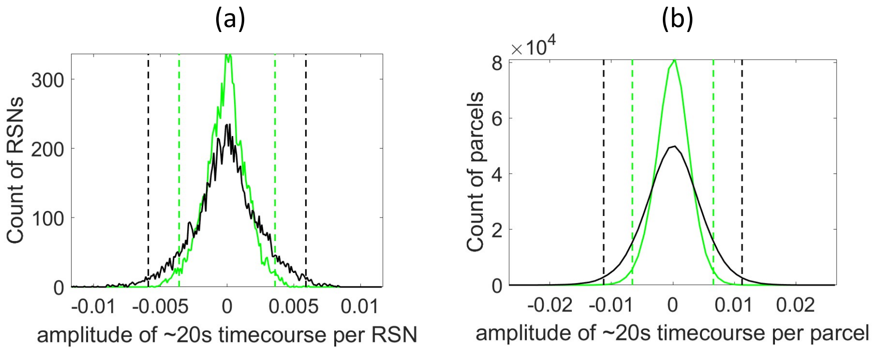


Fig.S14 Statistics to determine the magnitude threshold for activation and deactivation of a QPP's timecourse which is obtained by averaging timecourses of cortical vertices within each of Yeo's seven RSNs (a) or Glasser's 360 parcels (b). The process to determine these magnitude thresholds is similar to that used for active vertices/voxels, but the averages of the timecourses of vertices per null pattern are used instead, i.e., number of entries per histogram for each Ni random segments are 7x30x50 (a) and 360x30x50 (b). Magnitude threshold for all QPPs were set to 0.006 for a cortical RSN timecourse and 0.012 for a cortical parcel timecourse. For all QPPs, average timecourses across the cortical RSNs or the parcels chosen as the timing references (LPCC, V2, M1 and SMG) reach above the magnitude thresholds obtained here, showing they all entail significant activity.

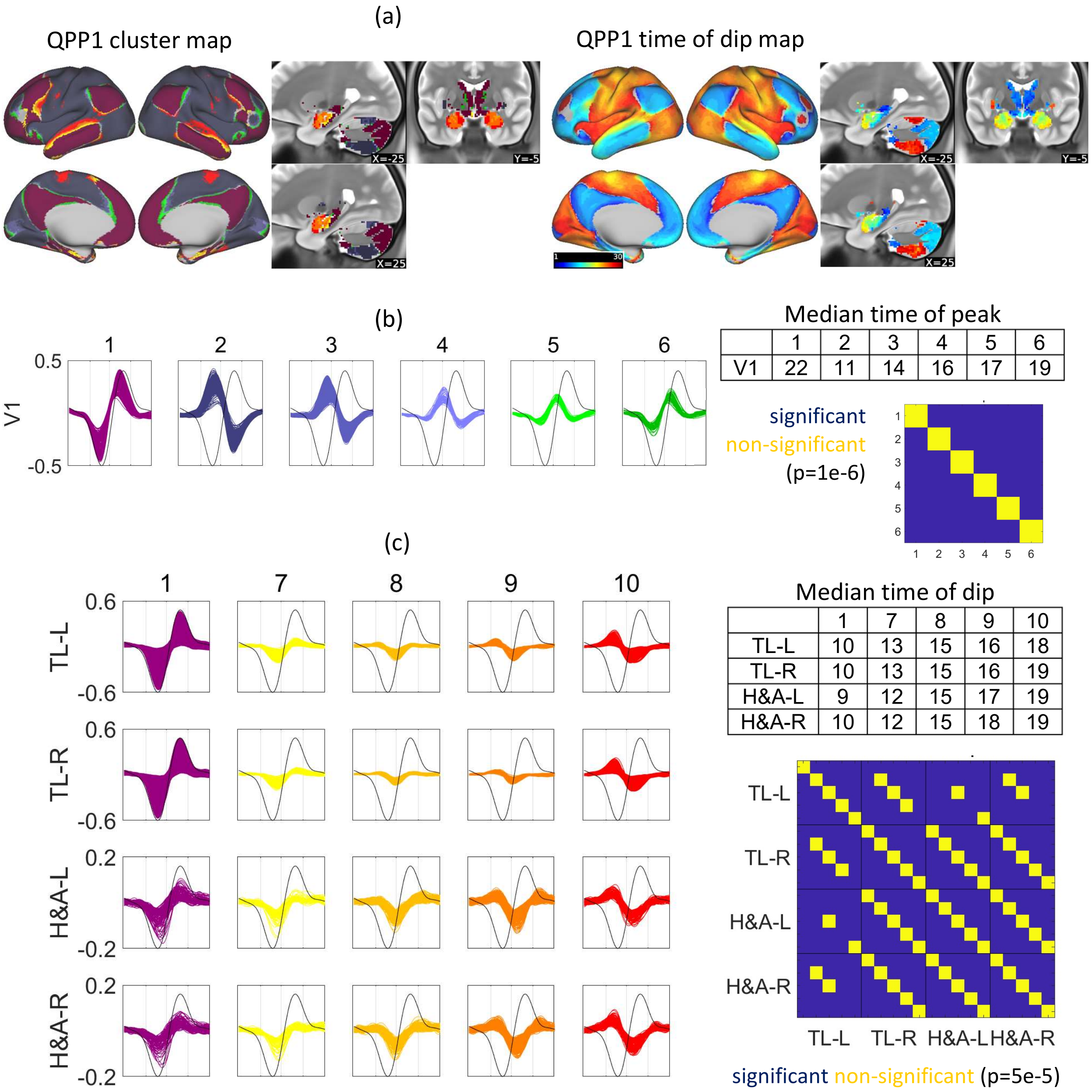


Median time of peak

| | 1 | 2 | 3 | 4 | 5 | 6 |
|---|---|---|---|---|---|---|
| V1 | 22 | 11 | 14 | 16 | 17 | 19 |

Median time of dip

| | 1 | 7 | 8 | 9 | 10 |
|---|---|---|---|---|---|
| TL-L | 10 | 13 | 15 | 16 | 18 |
| TL-R | 10 | 13 | 15 | 16 | 19 |
| H&A-L | 9 | 12 | 15 | 17 | 19 |
| H&A-R | 10 | 12 | 15 | 18 | 19 |

Fig.S15 Support for two focal propagations within QPP1 (for details see S.M.2, fourth paragraph). (a) Cluster map and time of dip map as spatial references for parts b and c. (b) Focal propagation within QPP1 across V1, supported by the order of the clusters of timecourses that tile V1 and the significant progression of their median peak times. (c) Focal propagations within QPP1 across the temporal Lobe (TL), and also across the hippocampus and amygdala (H&A), in the left (L) and right (R) hemispheres, supported by the order of the clusters of timecourses that tile TL and also tile H&A and the significant progression of their median dip times. Since the same clusters with the same order tile TL and H&A, with similar dip times (note the off diagonal entries of the significance matrix on the right, particularly TL-R vs H&A-R/-L), propagation of activity across TL (from medial to superior) and H&A happens at the same time.

Table S2 Size of QPP1's clusters per existing networks/parcels/probabilistic-maps per brain region (corresponding to S.M.3.3). For the cerebral cortex, cerebellum and striatum, such quantification supports the overall match between the activity within QPP1 and the adopted functional networks (Table S1). This match is mostly based on the first two clusters of QPP1, which are the largest and also are anticorrelated with one another. For other non-cortical regions, the quantification provided here mostly serves to identify which anatomical areas contain which clusters.

| | | **1** | **2** | **3** | **4** | **5** | **6** | **7** | **8** | **9** | **10** |
|---|---|---|---|---|---|---|---|---|---|---|---|
| **Cx** | **V** | 637 | 6642 | 628 | 104 | 123 | 229 | 16 | - | - | 151 |
| | **SM** | - | 10772 | 114 | - | - | - | - | - | - | 1074 |
| | **DA** | 220 | 5949 | 114 | 49 | 20 | 34 | 38 | - | 26 | 209 |
| | **VA** | 101 | 5987 | 509 | 147 | 140 | 163 | - | - | - | 42 |
| | **L** | 1994 | 598 | 78 | 41 | 35 | 72 | 277 | 140 | 101 | 372 |
| | **FP** | 5060 | 296 | 289 | 221 | 240 | 619 | 115 | 55 | 48 | 83 |
| | **DM** | 10923 | 328 | 25 | 17 | 55 | 143 | 420 | 182 | 134 | 394 |
| **C** | **V** | - | 25 | - | - | - | - | - | - | - | - |
| | **SM** | 348 | 326 | 82 | - | 17 | 41 | - | - | - | - |
| | **DA** | 84 | 955 | 146 | 31 | 23 | 25 | - | - | - | - |
| | **VA** | 166 | 1723 | 934 | 253 | 145 | 105 | - | - | - | - |
| | **L** | 467 | - | - | - | - | 20 | - | - | - | - |
| | **FP** | 3392 | 82 | 160 | 91 | 116 | 186 | - | - | - | - |
| | **DM** | 4323 | - | - | - | - | - | - | - | - | - |
| **T** | **PFC** | 205 | 127 | 40 | - | 116 | 309 | - | - | - | - |
| | **PreM** | - | 44 | - | - | - | 20 | - | - | - | - |
| | **M1** | - | 16 | - | - | - | - | - | - | - | - |
| | **S1** | - | 21 | - | - | - | - | - | - | - | - |
| | **PPC** | - | 163 | - | - | - | - | - | - | - | 18 |
| | **OCL** | - | - | - | - | - | - | - | - | - | - |
| | **TL** | 520 | - | - | - | - | 83 | - | - | - | - |
| **H** | **LA** | 17 | - | - | - | - | - | 26 | 50 | 35 | - |
| | **LM** | 41 | - | - | - | - | - | 29 | 33 | - | - |
| | **LP** | 28 | - | - | - | - | - | - | - | - | - |
| | **RA** | - | - | - | - | - | - | - | 15 | 22 | - |
| | **RAM** | - | - | - | - | - | - | - | 25 | 25 | - |
| | **RAL** | - | - | - | - | - | - | - | 33 | - | - |
| | **RM** | 19 | - | - | - | - | - | - | - | - | - |
| | **RP** | - | - | - | - | - | - | - | - | - | - |
| **A** | **P1** | - | - | - | - | - | - | - | - | 92 | 31 |
| | **P2** | - | - | - | - | - | - | - | 24 | 37 | - |
| | **P3** | - | - | - | - | - | - | - | - | 26 | - |
| | **P4** | - | - | - | - | - | - | - | - | 15 | 18 |
| | **P5** | - | - | - | - | - | - | - | - | 39 | 30 |
| | **P6** | - | - | - | - | - | - | - | - | 49 | 33 |
| **BD** | **LC** | 38 | - | - | - | - | - | - | - | - | - |
| | **VTA** | 29 | - | - | - | - | - | - | - | - | - |
| | **SNc** | 33 | - | - | - | - | 18 | - | - | - | - |
| | **SNr** | 109 | - | - | - | - | - | - | - | - | - |
| | **BNM** | 44 | - | - | - | - | - | - | - | - | - |
| | **PPN** | 46 | - | - | - | - | - | - | - | - | - |
| | **DR** | 74 | - | - | - | - | - | - | - | - | - |
| | **MR** | 40 | - | - | - | - | - | - | - | - | - |
| | **PAG** | 46 | - | - | - | - | - | - | - | - | - |
| | **STH** | - | - | - | - | - | - | - | - | - | - |
| | **GPe** | 15 | - | - | - | - | - | - | - | - | - |
| | **GPi** | - | - | - | - | - | - | - | - | - | - |
| | **HTH** | 170 | - | - | - | - | - | - | - | - | - |
| **S** | **V** | - | - | - | - | - | - | - | - | - | - |
| | **SM** | 52 | 40 | - | - | - | - | - | - | - | - |
| | **DA** | - | - | - | - | - | - | - | - | - | - |
| | **VA** | 21 | 25 | - | - | - | - | - | - | - | - |
| | **L** | 221 | - | - | - | - | 20 | - | - | - | - |
| | **FP** | 706 | - | - | - | - | 47 | - | - | - | - |
| | **DM** | 653 | - | - | - | - | 17 | - | - | - | - |

**Cerebral Cortex (Cx), Cerebellum (C) & Striatum (S)**
V: Visual
SM: SomatoMotor
DA: Dorsal Attention
VA: Ventral Attention
L: Limbic
FP: FrontoParietal
DM: Default Mode

**Thalamus (T)**
PFC: Prefrontal Cortex
PreM: PreMotor
M1: Primary Motor
S1: Primary Sensory
PPC: Posterior Parietal Cortex
OCL: Occipital Lobe
TL: Temporal Lobe

**Hippocampus (H)**
LA: Left Anterior
LM: Left Middle
LP: Left Posterior
RA: Right Anterior
RAM: Right Anteromedial
RAL: Right Anterolateral
RM: Right Medial
RP: Right Posterior

**Amygdala (A)**
P1 (Parcel1): lateral (La)
P2: basolateral (BL)
P3: accessory basal (BM)
P4: Paralaminar (BLV)
P5: corticomedial & central nuclei (CMN&CEN)
P6: ASTA&ATA&AAA&AMY

**Brainstem & Deep Brain Nuclei (BD)**
**LC:** Locus coeruleus
**VTA:** Ventral tegmental area
**SNc/SNr:** Substantia nigra compacta/reticulata
**BNM:** Basal nucleus of Meynert
**PPN:** Pedunculopontine nucleus
**Dr/MR:** Dorsal/Medial raphe
**PAG:** Periaqueductal gray
**STH:** subthalamic nucleus
**GPe/GPi:** Globus pallidus external/internal
**HTH:** Hypothalamus

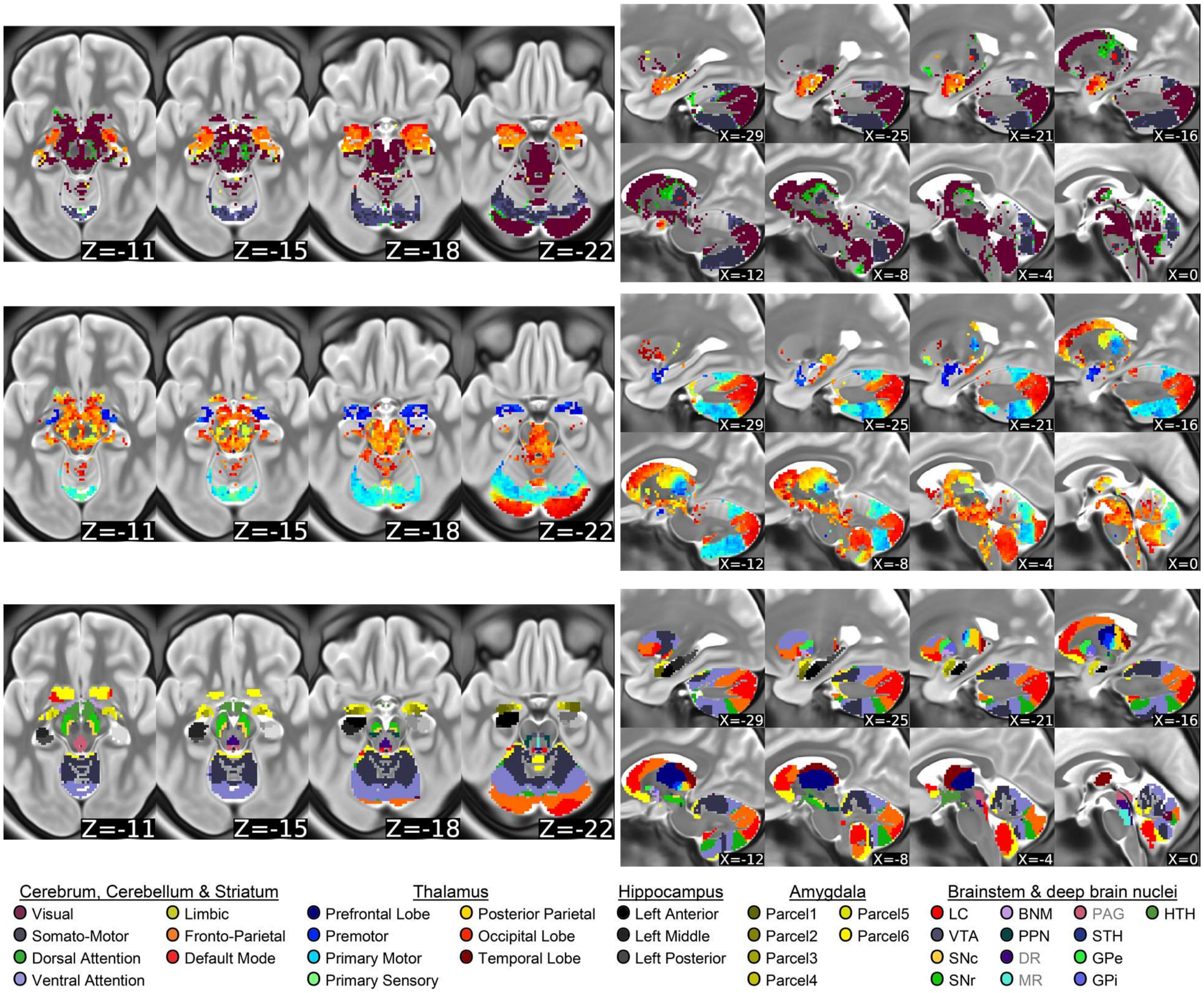


Fig.S16 Summary maps of QPP1 at additional planes along with the adopted parcellations for qualitative identification of which anatomical areas contain which clusters.

Cx: Cerebral cortex
C: Cerebellum
T: Thalamus
H: Hippocampus
A: Amygdala
BD: Brainstem and Deep brain
S: Striatum

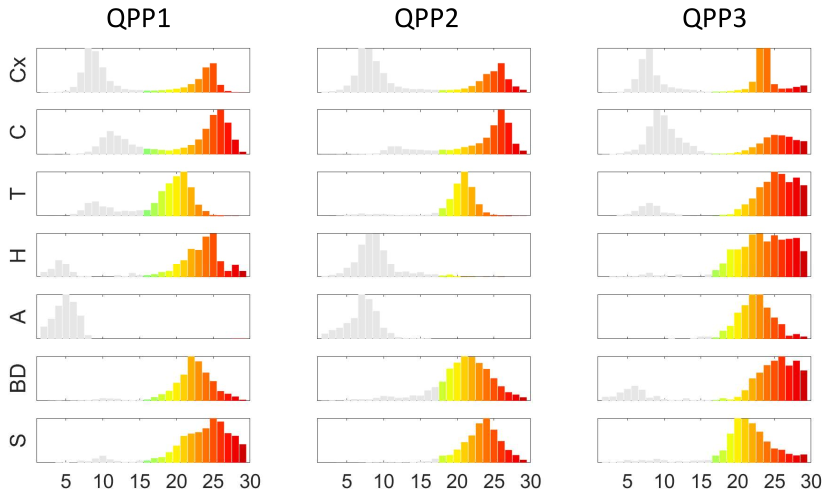


Median time of peak per region for the send half of QPP cycle

| | QPP1 | QPP2 | QPP3 |
|---|---|---|---|
| Cx | 24 | 25 | 24 |
| C | 25 | 26 | 26 |
| T | 20 | 21 | 25 |
| H | 24 | 19 | 24 |
| A | - | - | 22 |
| BD | 22 | 22 | 26 |
| S | 24 | 24 | 22 |

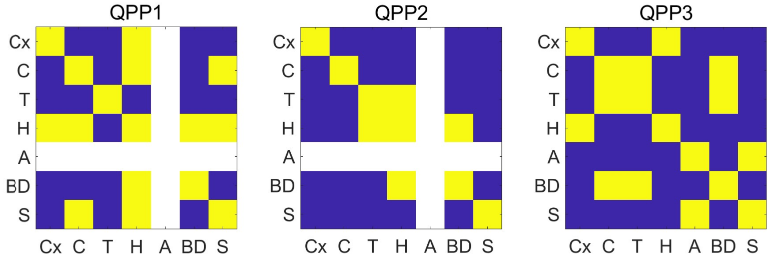


Fig.S17 Timing differences between brain regions that suggest driving mechanisms (for details about methods see S.M.4).

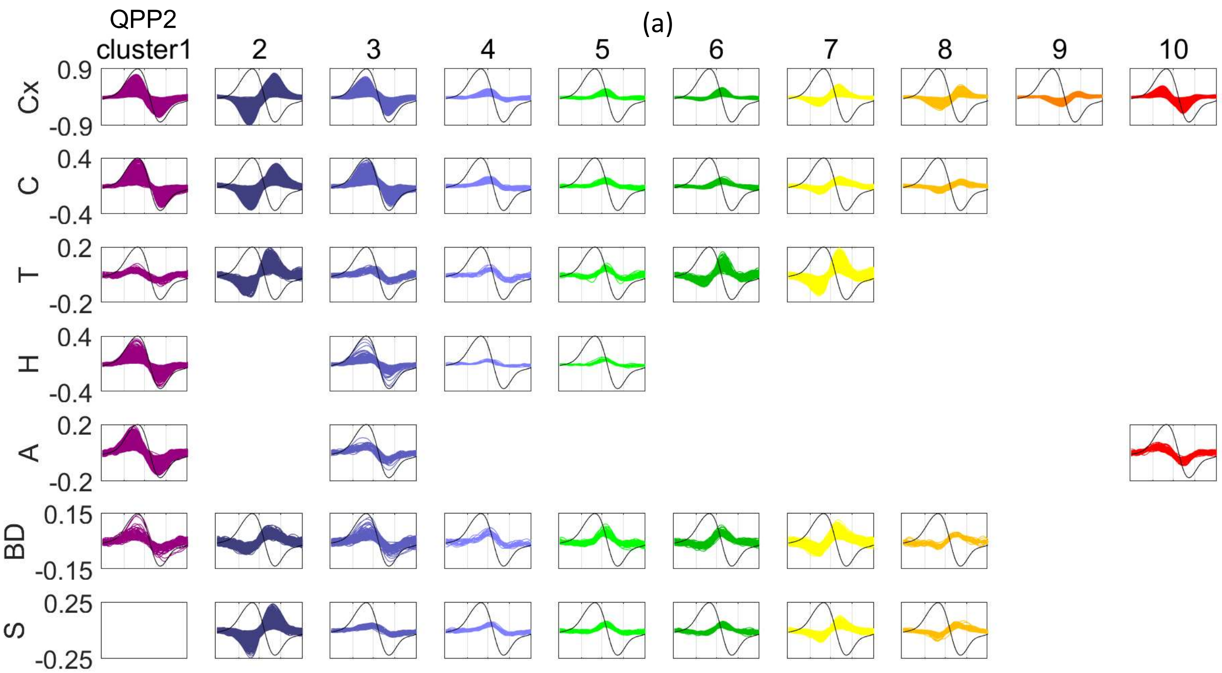


(b) Median time of peak

| | Cx | C | T | H | A | BD | S |
|---|---|---|---|---|---|---|---|
| **1** | 8 | 10 | 8 | 8 | 7 | 9 | - |
| **2** | 25 | 26 | 22 | - | - | 23 | 24 |
| **3** | 12 | 12 | 13 | 11 | 10 | 13 | 15 |
| **4** | 16 | 16 | 17 | 15 | - | 16 | 17 |
| **5** | 17 | 17 | 18 | 17 | - | 17 | 18 |
| **6** | 18 | 18 | 19 | - | - | 19 | 19 |
| **7** | 20 | 21 | 21 | - | - | 21 | 21 |
| **8** | 27 | 25 | - | - | - | 23 | 23 |
| **9** | 27 | - | - | - | - | - | - |
| **10** | 6 | - | - | - | 6 | - | - |

(c) Significance of difference in time of peak
significant non-significant (p=4e-6)

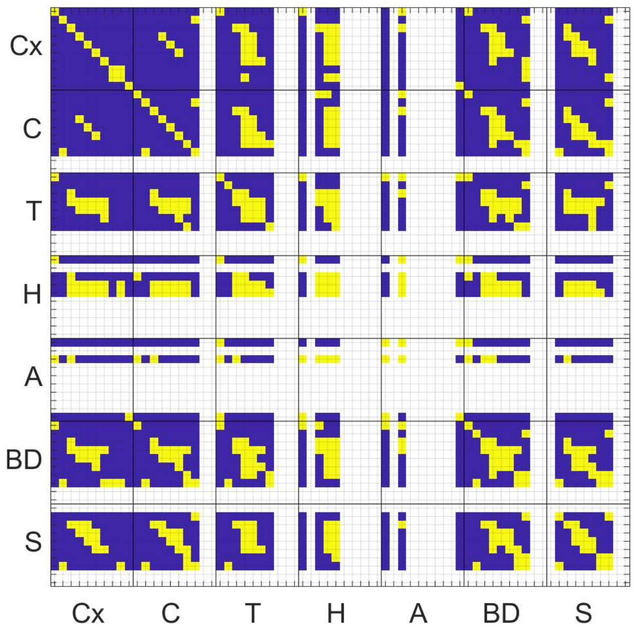


(d) Cluster size

| | Cx | C | T | H | A | BD | S |
|---|---|---|---|---|---|---|---|
| **1** | 26445 | 486 | 128 | 1053 | 484 | 71 | - |
| **2** | 16627 | 9506 | 521 | - | - | 885 | 2644 |
| **3** | 4499 | 1763 | 73 | 282 | 45 | 106 | 56 |
| **4** | 517 | 248 | 21 | 16 | - | 20 | 45 |
| **5** | 519 | 232 | 28 | 16 | - | 55 | 55 |
| **6** | 425 | 225 | 136 | - | - | 97 | 62 |
| **7** | 1224 | 905 | 780 | - | - | 658 | 445 |
| **8** | 1997 | 314 | - | - | - | 25 | 52 |
| **9** | 913 | - | - | - | - | - | - |
| **10** | 2476 | - | - | - | 76 | - | - |

Fig.S18 Support for the propagation of activity and coactivity between non-adjacent areas within QPP2 (see S.M.2 for details). (a) QPP2's clusters of timecourses per region (see Fig.2e for the map of clusters; LPCC in black for reference). (b) Median time of peak per cluster per region. (c) Significance of differences in the time of peak between included clusters using dependent t-test (p=0.01/(70x69/2)). (d) Cluster size per region. The propagation of activity per region within QPP2 is supported by the significant progression of the median time-of-peak across clusters timecourses taken together with the spatial order of these clusters. The coactivity of non-adjacent areas (e.g., non-cortical and cortical areas in Table 1) is supported by those areas belonging to the same cluster, despite slight but significant time differences.

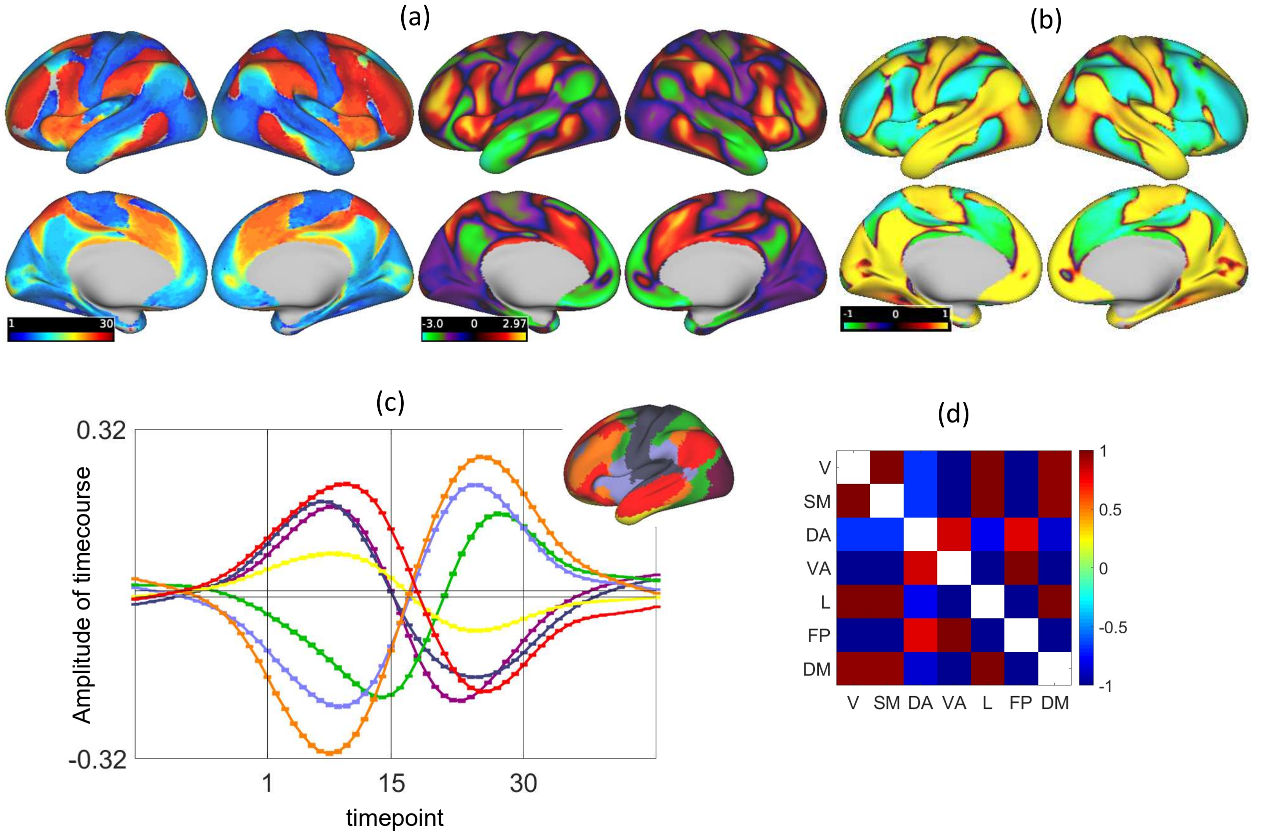


(e)

| | Time of peak | Time of zero-crossing | Number of active vertices |
|---|---|---|---|
| V | 9 | 15 | 8781 |
| SM | 7 | 15 | 11909 |
| DA | 26 | 18 | 6752 |
| VA | 24 | 16 | 7089 |
| L | 9 | 17 | 3885 |
| FP | 26 | 17 | 7264 |
| DM | 10 | 17 | 12772 |

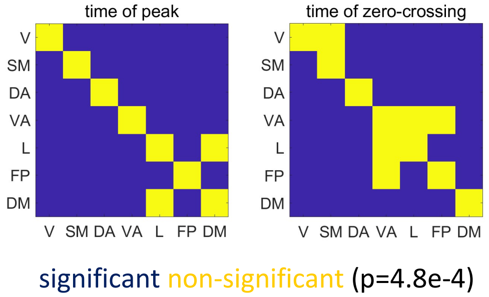


Fig.S19 (a) QPP2's time of peak map versus FCG3. The correlation between these two maps is 0.86, supporting that the peak activity within QPP2 sweeps FCG3. (b) Correlation map within QPP2, with LPCC being the seed timecourse, is slightly more similar to FCG3 compared to the time of peak map (correlation: -0.91). (c) Seven timecourses, each the average of the QPP2's timecourses across a cortical RSN (map of RSNs in the top right serves as the legend; all timecourses entail significant activity, see Fig.S14a; error bars show the standard error). (d) Correlation between timecourses shown in (c). (e) Median of time of peak and time of zero-crossing per group (i.e., RSN). For details about the methods, see S.M.3.1 and SM.3.2.

Table S3 Size of QPP2's clusters per existing networks/parcels/probabilistic-maps per region (corresponding to S.M.3.3). For the cerebral cortex, cerebellum and striatum, such quantification supports the overall match between the activity within QPP2 and the adopted functional networks. This match is mostly based on the first two clusters of QPP2, which are the largest and also are anticorrelated with one another. For other non-cortical regions, the quantification provided here mostly serves to identify which anatomical areas contain which clusters.

| | | 1 | 2 | 3 | 4 | 5 | 6 | 7 | 8 | 9 | 10 |
|---|---|---|---|---|---|---|---|---|---|---|---|
| Cx | V | 5513 | 210 | 1247 | 73 | 136 | 84 | 215 | 62 | 58 | 810 |
| | SM | 9677 | 142 | 472 | 108 | 76 | 35 | 39 | 112 | 53 | 270 |
| | DA | 595 | 2785 | - | - | - | - | - | 1209 | 624 | 970 |
| | VA | 406 | 5483 | 16 | - | - | 35 | 160 | 422 | 102 | 110 |
| | L | 2430 | 714 | 320 | - | - | - | 85 | 42 | - | 162 |
| | FP | 67 | 6477 | 38 | - | 18 | 17 | 191 | 121 | 50 | 66 |
| | DM | 7757 | 816 | 2405 | 314 | 281 | 241 | 534 | 29 | 15 | 88 |
| C | V | - | - | - | - | - | - | - | - | - | - |
| | SM | 24 | 374 | 19 | - | - | 22 | 171 | 26 | - | - |
| | DA | - | 1063 | - | - | - | - | 18 | 120 | - | - |
| | VA | 16 | 2785 | - | - | - | - | 142 | 138 | - | - |
| | L | - | 262 | 65 | - | - | 19 | 102 | - | - | - |
| | FP | - | 4137 | - | - | - | - | 54 | 20 | - | - |
| | DM | 433 | 875 | 1669 | 232 | 209 | 172 | 414 | - | - | - |
| T | PFC | 17 | 317 | 19 | - | - | 65 | 442 | - | - | - |
| | PreM | - | - | - | - | - | - | 42 | - | - | - |
| | M1 | - | - | - | - | - | - | - | - | - | - |
| | S1 | 20 | - | - | - | - | - | - | - | - | - |
| | PPC | 44 | 29 | - | - | - | - | 17 | - | - | - |
| | OCL | - | - | - | - | - | - | - | - | - | - |
| | TL | - | 155 | 42 | 15 | 17 | 63 | 264 | - | - | - |
| H | LA | 155 | - | - | - | - | - | - | - | - | - |
| | LM | 118 | - | 15 | - | - | - | - | - | - | - |
| | LP | 48 | - | 33 | - | - | - | - | - | - | - |
| | RA | 70 | - | - | - | - | - | - | - | - | - |
| | RAM | 59 | - | - | - | - | - | - | - | - | - |
| | RAL | 86 | - | - | - | - | - | - | - | - | - |
| | RM | 64 | - | 22 | - | - | - | - | - | - | - |
| | RP | 22 | - | 22 | - | - | - | - | - | - | - |
| A | P1 | 104 | - | - | - | - | - | - | - | - | - |
| | P2 | 38 | - | - | - | - | - | - | - | - | 16 |
| | P3 | 37 | - | - | - | - | - | - | - | - | - |
| | P4 | 24 | - | - | - | - | - | - | - | - | - |
| | P5 | 68 | - | - | - | - | - | - | - | - | - |
| | P6 | 54 | - | - | - | - | - | - | - | - | - |
| BD | LC | - | - | - | - | - | - | - | - | - | - |
| | VTA | - | - | - | - | - | - | - | - | - | - |
| | SNc | - | 38 | - | - | - | - | 21 | - | - | - |
| | SNr | - | 60 | - | - | - | - | 36 | - | - | - |
| | BNM | - | 20 | - | - | - | - | 24 | - | - | - |
| | PPN | - | 20 | - | - | - | - | - | - | - | - |
| | DR | - | - | - | - | - | - | - | - | - | - |
| | MR | - | - | - | - | - | - | - | - | - | - |
| | PAG | - | 23 | - | - | - | - | 17 | - | - | - |
| | STH | - | - | - | - | - | - | - | - | - | - |
| | GPe | - | 115 | - | - | - | - | 50 | - | - | - |
| | GPi | - | 33 | - | - | - | - | 45 | - | - | - |
| | HTH | 30 | - | 35 | - | - | - | 21 | - | - | - |
| S | V | - | - | - | - | - | - | - | - | - | - |
| | SM | - | 320 | - | - | - | - | - | 20 | - | - |
| | DA | - | 39 | - | - | - | - | - | - | - | - |
| | VA | - | 648 | - | - | - | - | 18 | - | - | - |
| | L | - | 69 | 23 | 21 | 24 | 23 | 84 | - | - | - |
| | FP | - | 1051 | - | - | - | - | 123 | - | - | - |
| | DM | - | 406 | 24 | 20 | 29 | 37 | 189 | - | - | - |

**Cerebral Cortex (Cx), Cerebellum (C) & Striatum (S)**
V: Visual
SM: SomatoMotor
DA: Dorsal Attention
VA: Ventral Attention
L: Limbic
FP: FrontoParietal
DM: Default Mode

**Thalamus (T)**
PFC: Prefrontal Cortex
PreM: PreMotor
M1: Primary Motor
S1: Primary Sensory
PPC: Posterior Parietal Cortex
OCL: Occipital Lobe
TL: Temporal Lobe

**Hippocampus (H)**
LA: Left Anterior
LM: Left Middle
LP: Left Posterior
RA: Right Anterior
RAM: Right Anteromedial
RAL: Right Anterolateral
RM: Right Medial
RP: Right Posterior

**Amygdala (A)**
P1 (Parcel1): lateral (La)
P2: basolateral (BL)
P3: accessory basal (BM)
P4: Paralaminar (BLV)
P5: corticomedial & central nuclei (CMN&CEN)
P6: ASTA&ATA&AAA&AMY

**Brainstem & Deep Brain Nuclei (BD)**
**LC:** Locus coeruleus
**VTA:** Ventral tegmental area
**SNc/SNr:** Substantia nigra compacta/reticulata
**BNM:** Basal nucleus of Meynert
**PPN:** Pedunculopontine nucleus
**Dr/MR:** Dorsal/Medial raphe
**PAG:** Periaqueductal gray
**STH:** subthalamic nucleus
**GPe/GPi:** Globus pallidus external/internal
**HTH:** Hypothalamus

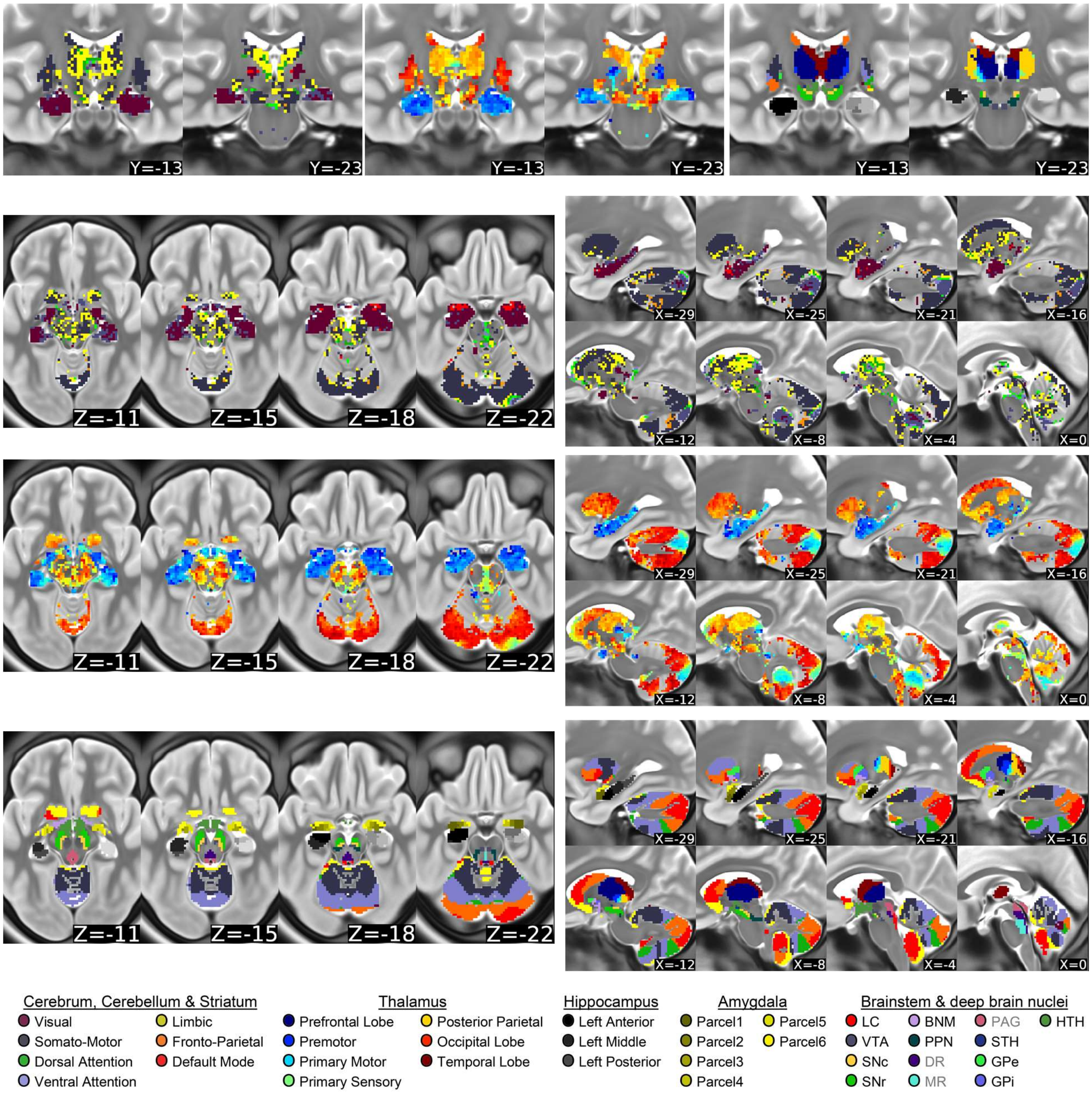


Fig.S20 Summary maps of QPP2 at additional planes along with the adopted parcellations for qualitative identification of which anatomical areas contain which clusters.

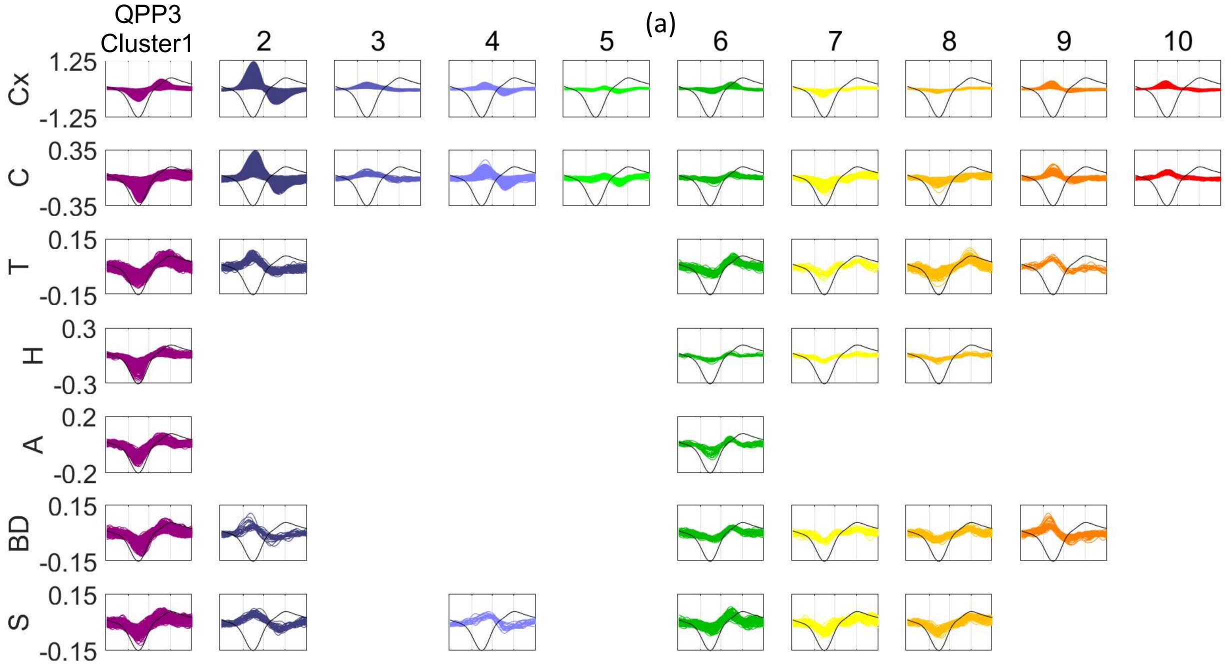


(b) Median time of peak

| | Cx | C | T | H | A | BD | S |
|---|---|---|---|---|---|---|---|
| **1** | 24 | 26 | 25 | 24 | 23 | 26 | 23 |
| **2** | 8 | 9 | 8 | - | - | 7 | 9 |
| **3** | 8 | 8 | - | - | - | - | - |
| **4** | 12 | 12 | - | - | - | - | 12 |
| **5** | 14 | 14 | - | - | - | - | - |
| **6** | 23 | 24 | 23 | - | 22 | 23 | 22 |
| **7** | 18.5 | 29 | - | - | - | - | 28 |
| **8** | 29 | 28 | 28 | - | - | 28 | - |
| **9** | 6 | 8 | 7 | - | - | 5.5 | - |
| **10** | 7 | 8 | - | - | - | - | - |

(c) Significance of difference in time of peak
significant non-significant (p=4e-6)

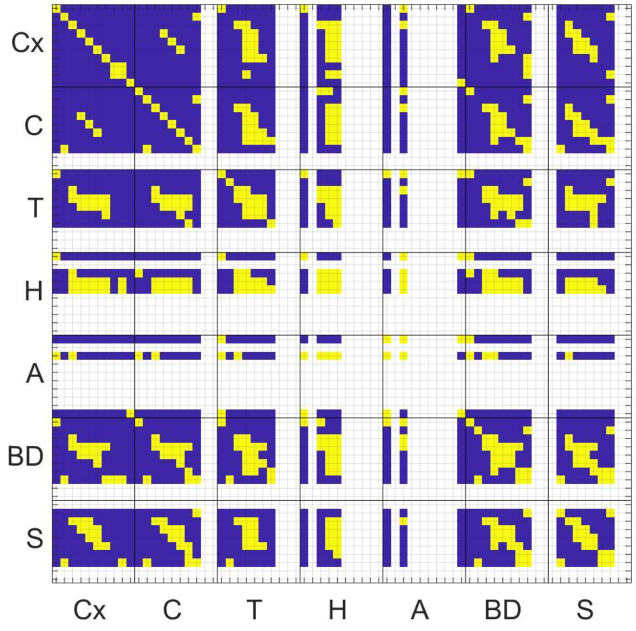


(d) Cluster size

| | Cx | C | T | H | A | BD | S |
|---|---|---|---|---|---|---|---|
| **1** | 25574 | 5458 | 874 | 838 | 282 | 646 | 978 |
| **2** | 18607 | 4011 | 113 | - | - | 28 | 41 |
| **3** | 519 | 136 | - | - | - | - | - |
| **4** | 1997 | 1559 | - | - | - | - | 19 |
| **5** | 395 | 221 | - | - | - | - | - |
| **6** | 2151 | 261 | 82 | 20 | 33 | 79 | 197 |
| **7** | 1141 | 777 | 23 | 51 | - | 36 | 92 |
| **8** | 285 | 331 | 110 | 38 | - | 69 | 54 |
| **9** | 1460 | 84 | 15 | - | - | 57 | - |
| **10** | 585 | 93 | - | - | - | - | - |

Fig.S21 Support for coactivity between non-adjacent areas and the propagation of activity within QPP3 (see S.M.2 for details). (a) QPP3's clusters of timecourses per region (see Fig.2f for the map of clusters; LPCC in black for reference). (b) Median time of peak per cluster per region. (c) Significance of differences in the time of peak between included clusters using dependent t-test (p=0.01/(70x69/2)). (d) Cluster size per region. The coactivity of non-adjacent areas (e.g., non-cortical and cortical areas in Table 1) is supported by those areas belonging to the same cluster .The propagation of activity is supported by the significant progression of the median time-of-peak across clusters timecourses taken together with the spatial order of these clusters.

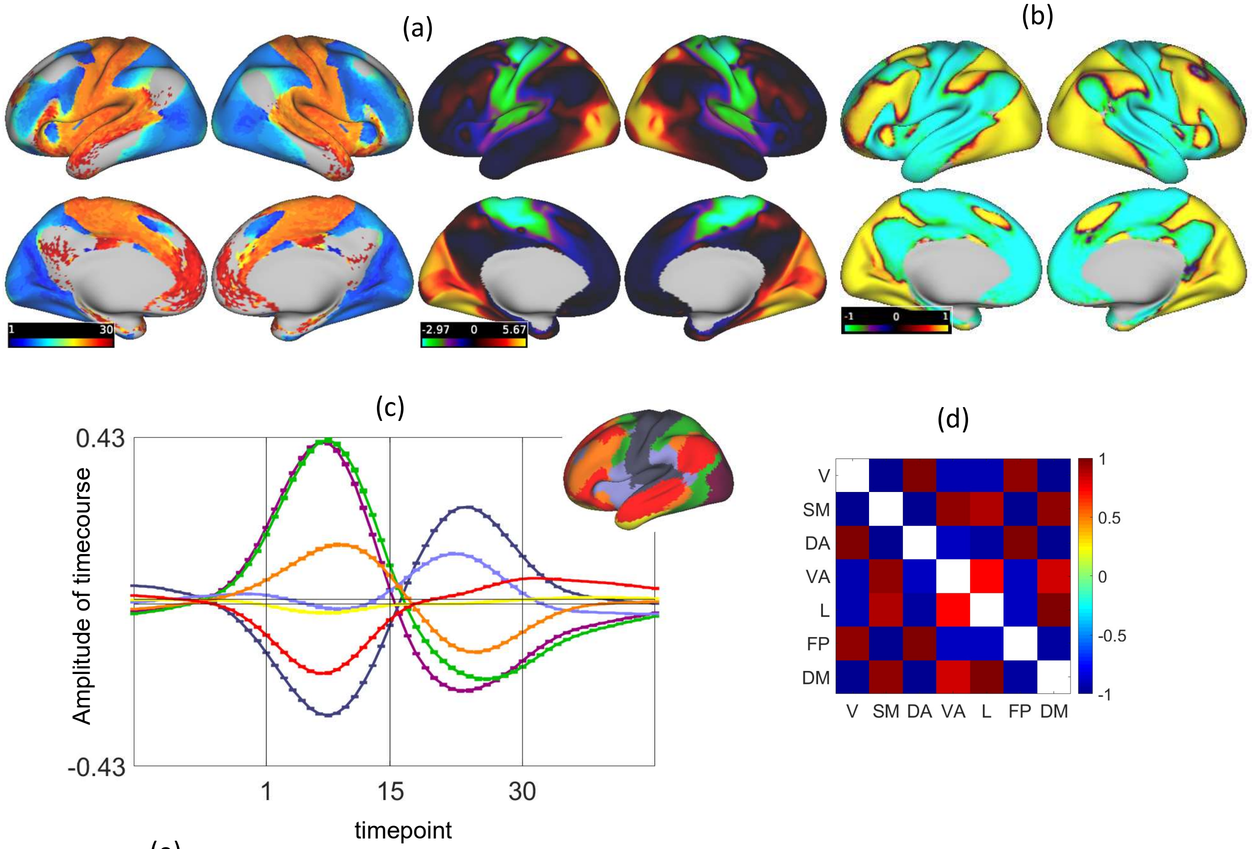


(e)

| | Time of peak | Time of zero-crossing | Number of active vertices |
|---|---|---|---|
| V | 7 | 15 | 8740 |
| SM | 24 | 16 | 11960 |
| DA | 8 | 16 | 6761 |
| VA | 22 | 15 | 7156 |
| L | 11 | 17 | 3537 |
| FP | 10 | 17 | 7237 |
| DM | 25 | 18 | 12697 |

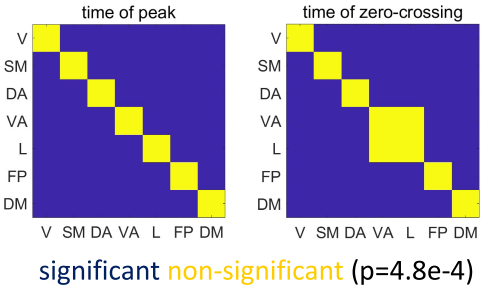


Fig.S22 (a) QPP3's time of peak map versus FCG2. The correlation between these two maps is -0.73, supporting that the peak activity within QPP3 matches FCG3. (b) Correlation map within QPP3, with V2 being the seed timecourse, is also similar to FCG3 (correlation: 0.72). (c) Seven timecourses, each the average of the QPP3's timecourses across a cortical RSN (map of RSNs in the top right serves as the legend; all timecourses entail significant activity, see Fig.S14a; error bars show the standard error). (d) Correlation between timecourses shown in (c). (e) Median of time of peak and time of zero-crossing per group (i.e., RSN). For details about the methods, see S.M.3.1 and SM.3.2.

Table S4 Size of QPP3's clusters per existing networks/parcels/probabilistic-maps per region (corresponding to S.M.3.3). For the cerebral cortex, cerebellum and striatum, such quantification supports the overall match between the activity within QPP3 and the adopted functional networks. This match is mostly based on the first two clusters of QPP3, which are the largest and also are anticorrelated with one another. For other non-cortical regions, the quantification provided here mostly serves to identify which anatomical areas contain which clusters.

| | | 1 | 2 | 3 | 4 | 5 | 6 | 7 | 8 | 9 | 10 |
|---|---|---|---|---|---|---|---|---|---|---|---|
| | **V** | 27 | 6953 | - | 229 | 73 | - | - | 42 | 1294 | - |
| | **SM** | 11424 | - | - | - | - | 378 | - | - | - | - |
| | **DA** | 297 | 5405 | 25 | - | - | 184 | - | - | - | 344 |
| **Cx** | **VA** | 2640 | 671 | 261 | 25 | - | 1514 | - | - | 45 | 179 |
| | **L** | 1646 | 1075 | 45 | 142 | - | 33 | 100 | 53 | 34 | 25 |
| | **FP** | 570 | 4136 | 180 | 1210 | 178 | 23 | 59 | - | - | 23 |
| | **DM** | 8970 | 367 | - | 390 | 138 | 18 | 981 | 189 | 78 | - |
| | **V** | - | 24 | - | - | - | - | - | - | - | - |
| | **SM** | 2166 | 25 | - | - | - | 94 | 43 | 91 | - | - |
| | **DA** | 84 | 979 | 20 | 43 | - | - | - | - | 15 | 27 |
| **C** | **VA** | 1241 | 814 | 71 | 146 | - | 143 | - | 42 | 45 | 56 |
| | **L** | 236 | 87 | - | - | - | - | 33 | 44 | - | - |
| | **FP** | 47 | 1934 | 39 | 1114 | 129 | - | 34 | - | - | - |
| | **DM** | 1681 | 148 | - | 239 | 90 | - | 648 | 146 | - | - |
| | **PFC** | 336 | - | - | - | - | 32 | - | 50 | - | - |
| | **PreM** | 102 | - | - | - | - | - | - | - | - | - |
| | **M1** | 44 | - | - | - | - | - | - | - | - | - |
| **T** | **S1** | 38 | - | - | - | - | - | - | - | - | - |
| | **PPC** | 89 | 31 | - | - | - | - | - | - | - | - |
| | **OCL** | - | 24 | - | - | - | - | - | - | - | - |
| | **TL** | 254 | 58 | - | - | - | - | 17 | 58 | - | - |
| | **LA** | 124 | - | - | - | - | - | - | - | - | - |
| | **LM** | 103 | - | - | - | - | - | - | - | - | - |
| | **LP** | 45 | - | - | - | - | - | - | - | - | - |
| **H** | **RA** | 63 | - | - | - | - | - | - | - | - | - |
| | **RAM** | 50 | - | - | - | - | - | - | - | - | - |
| | **RAL** | 76 | - | - | - | - | - | - | - | - | - |
| | **RM** | 41 | - | - | - | - | - | - | - | - | - |
| | **RP** | 15 | - | - | - | - | - | - | - | - | - |
| | **P1** | 63 | - | - | - | - | - | - | - | - | - |
| | **P2** | 17 | - | - | - | - | - | - | - | - | - |
| **A** | **P3** | 27 | - | - | - | - | - | - | - | - | - |
| | **P4** | - | - | - | - | - | - | - | - | - | - |
| | **P5** | 60 | - | - | - | - | - | - | - | - | - |
| | **P6** | 28 | - | - | - | - | - | - | - | - | - |
| | **LC** | - | - | - | - | - | - | - | - | - | - |
| | **VTA** | - | - | - | - | - | - | - | - | - | - |
| | **SNc** | - | - | - | - | - | - | - | - | - | - |
| | **SNr** | - | - | - | - | - | - | - | - | - | - |
| | **BNM** | 47 | - | - | - | - | - | - | - | - | - |
| | **PPN** | - | - | - | - | - | - | - | - | - | - |
| **BD** | **DR** | - | - | - | - | - | - | - | - | - | - |
| | **MR** | - | - | - | - | - | - | - | - | - | - |
| | **PAG** | - | - | - | - | - | - | - | - | - | - |
| | **STH** | - | - | - | - | - | - | - | - | - | - |
| | **GPe** | 43 | - | - | - | - | 22 | - | - | - | - |
| | **GPi** | 31 | - | - | - | - | - | - | - | - | - |
| | **HTH** | 54 | - | - | - | - | - | - | - | - | - |
| | **V** | - | - | - | - | - | - | - | - | - | - |
| | **SM** | 277 | - | - | - | - | 46 | - | - | - | - |
| | **DA** | - | - | - | - | - | - | - | - | - | - |
| **S** | **VA** | 56 | - | - | - | - | 45 | - | - | - | - |
| | **L** | 149 | - | - | - | - | - | 15 | - | - | - |
| | **FP** | 128 | 31 | - | - | - | 34 | - | - | - | - |
| | **DM** | 287 | - | - | - | - | 44 | 61 | 33 | - | - |

**Cerebral Cortex (Cx), Cerebellum (C) & Striatum (S)**
V: Visual
SM: SomatoMotor
DA: Dorsal Attention
VA: Ventral Attention
L: Limbic
FP: FrontoParietal
DM: Default Mode

**Thalamus (T)**
PFC: Prefrontal Cortex
PreM: PreMotor
M1: Primary Motor
S1: Primary Sensory
PPC: Posterior Parietal Cortex
OCL: Occipital Lobe
TL: Temporal Lobe

**Hippocampus (H)**
LA: Left Anterior
LM: Left Middle
LP: Left Posterior
RA: Right Anterior
RAM: Right Anteromedial
RAL: Right Anterolateral
RM: Right Medial
RP: Right Posterior

**Amygdala (A)**
P1 (Parcel1): lateral (La)
P2: basolateral (BL)
P3: accessory basal (BM)
P4: Paralaminar (BLV)
P5: corticomedial & central nuclei (CMN&CEN)
P6: ASTA&ATA&AAA&AMY

**Brainstem & Deep Brain Nuclei (BD)**
**LC:** Locus coeruleus
**VTA:** Ventral tegmental area
**SNc/SNr:** Substantia nigra compacta/reticulata
**BNM:** Basal nucleus of Meynert
**PPN:** Pedunculopontine nucleus
**Dr/MR:** Dorsal/Medial raphe
**PAG:** Periaqueductal gray
**STH:** subthalamic nucleus
**GPe/GPi:** Globus pallidus external/internal
**HTH:** Hypothalamus

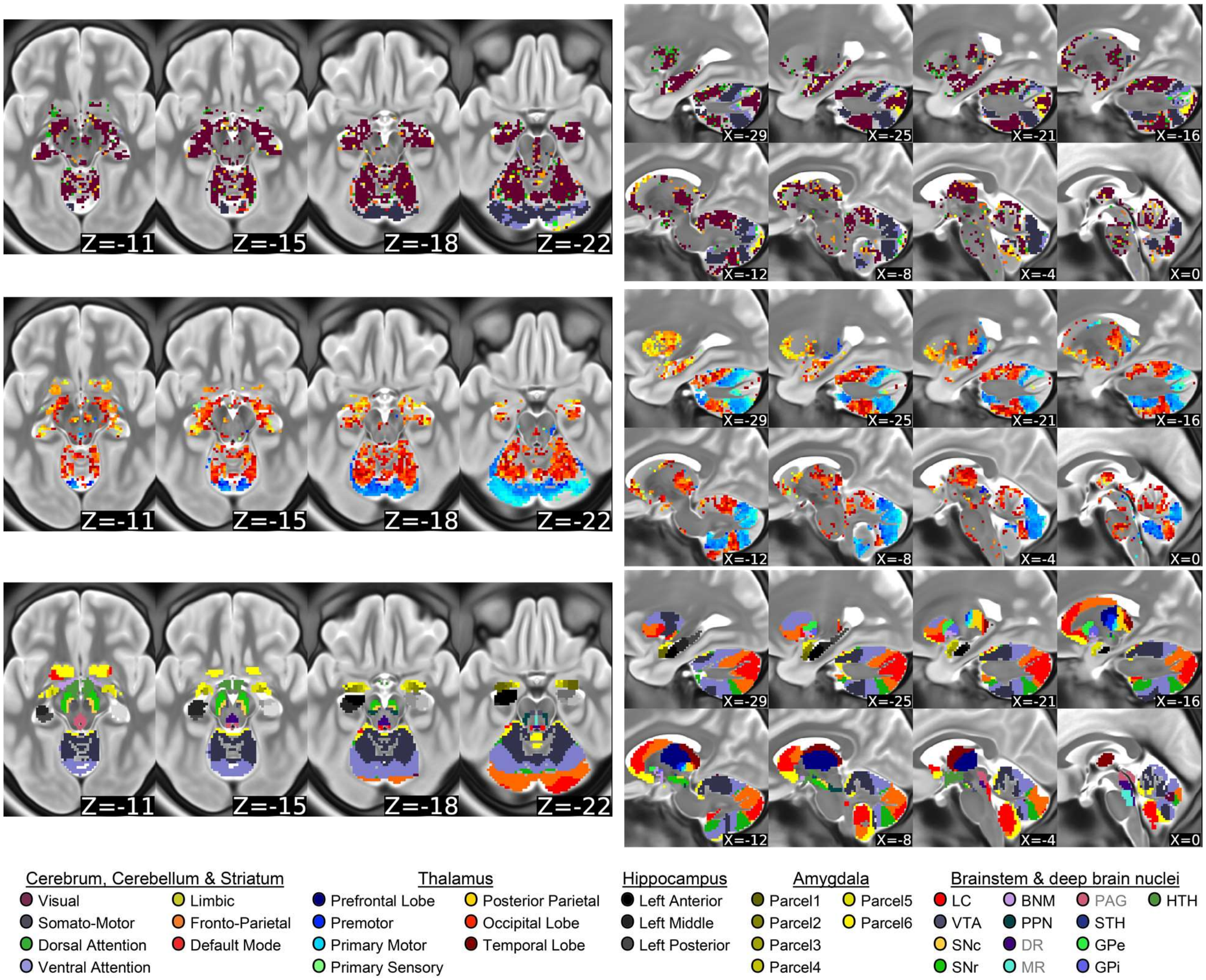


Fig.S23 Summary maps of QPP3 at additional planes along with the adopted parcellations for qualitative identification of which anatomical areas contain which clusters.

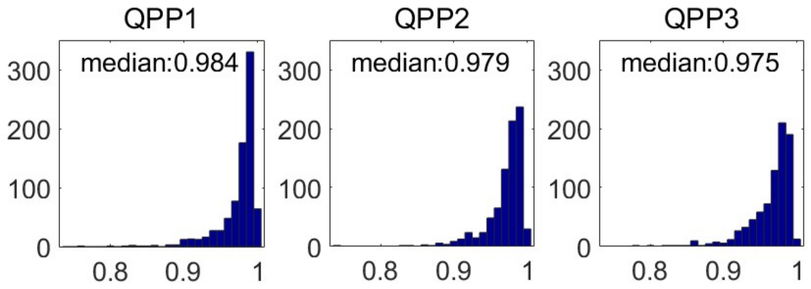


Fig.S24 Histograms and vertical axes show correlation between each QPP of each individual (detected and built based on WM-CSF-GM-regressed timeseries, i.e., GS-regressed timeseries) and the average of its contributing segments over the WM-CSF-regressed timeseries. Near identical outcome proves GSR does not influence QPPs 1-3GS (WM: white-matter, GM: gray matter, GS: global signal).

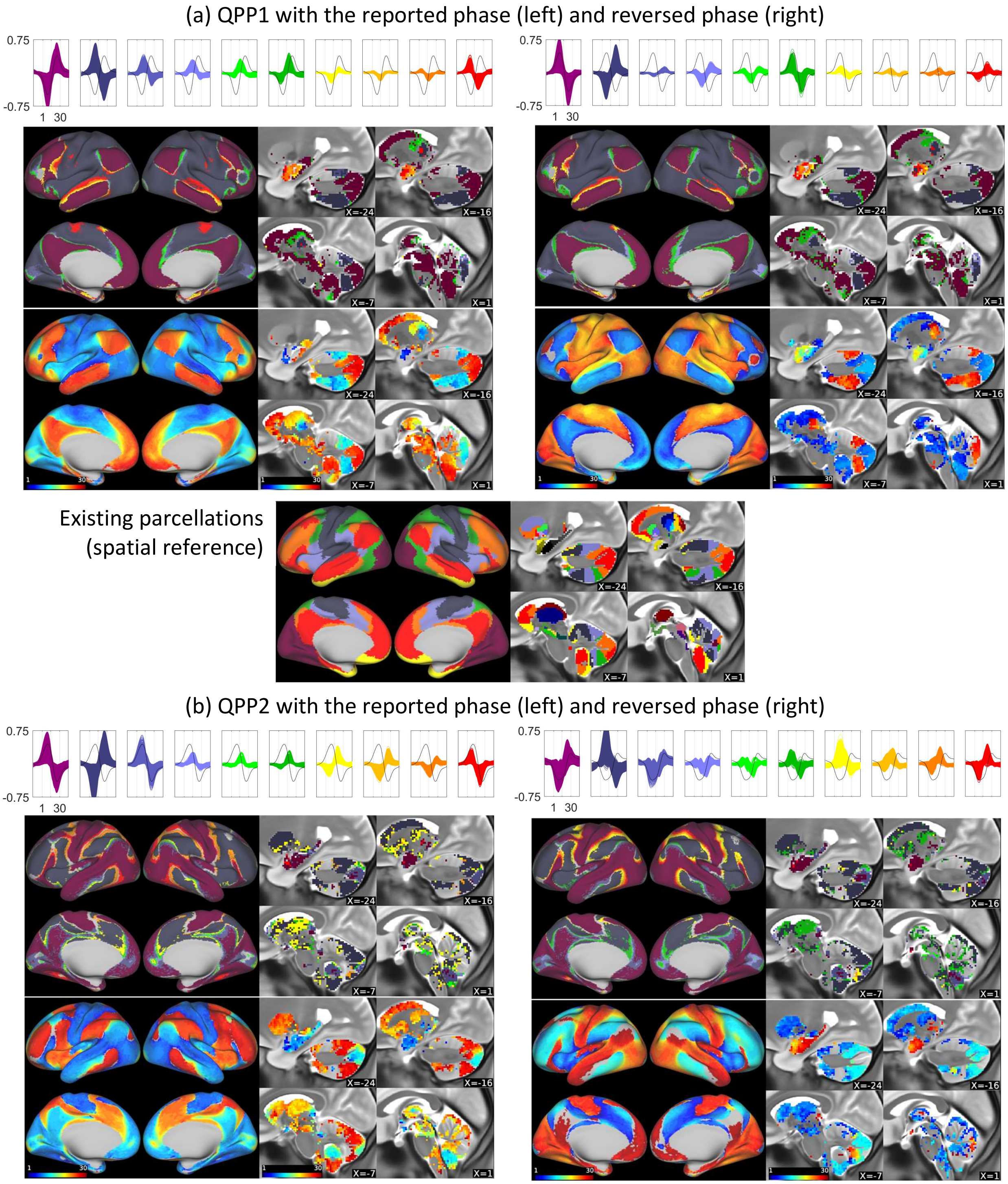


Fig.S25 QPPs with reversed phase, which have the main characteristics described in the main text, showing the phase of QPP, set by the choice of the seed parcel in the phase-adjustment step does not influence our results and conclusion. Note in QPP3 with reversed phase, the focal propagation does not exactly reverses (compare Table 1-2 and Video 3) and is as follows: SM→VA→DA/Vlateral & SM→DA/Vlateral & Vmedial-periphery→Vmedial-fovea/FPpostero-medial & VA→FPlateral

(c) QPP3 with the reported phase (left) and reversed phase (right)

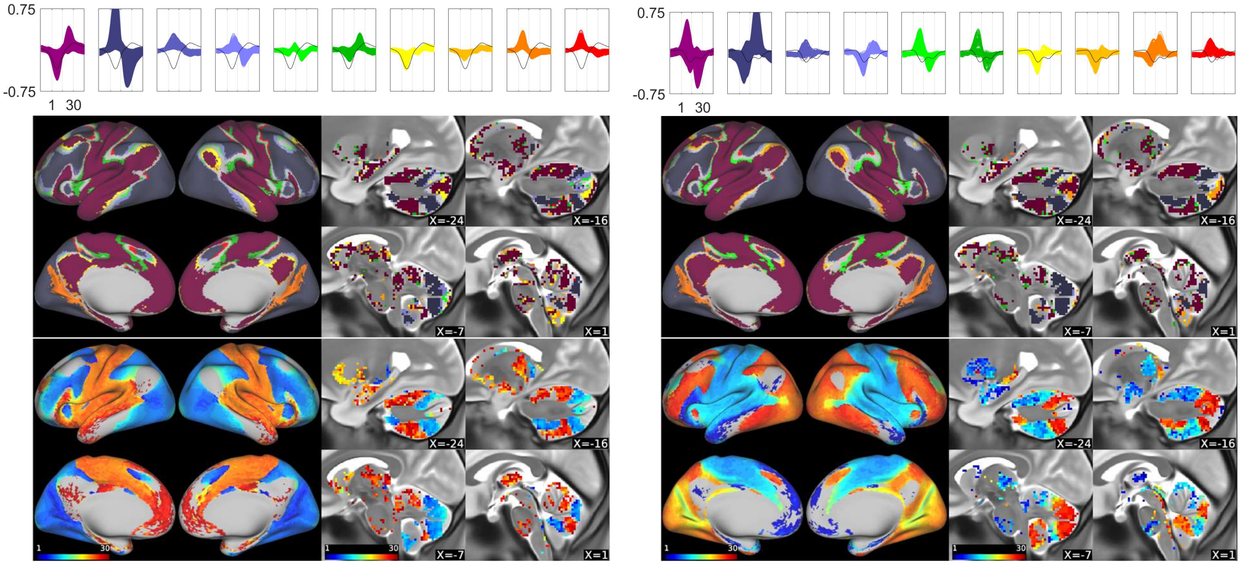


(d) Histograms of time of peak per region with the reported (left) and reversed (right) phase

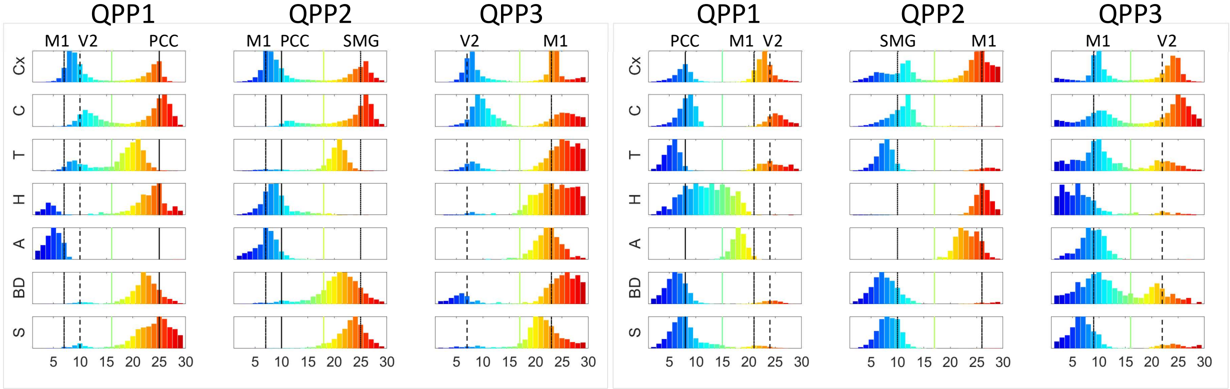


Significance of time of peak differences for the second half of QPP cycle

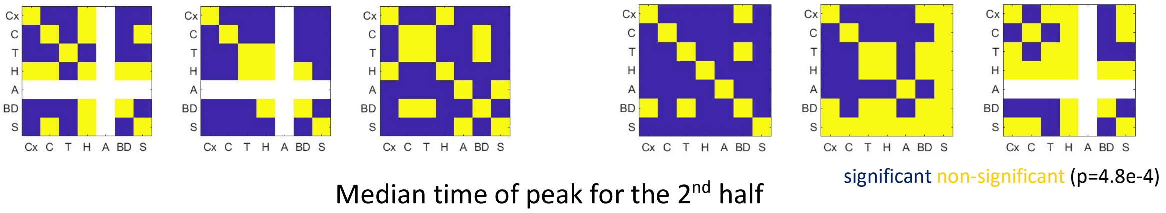


Median time of peak for the 2$^{nd}$ half

| | QPP1 | QPP2 | QPP3 | | QPP1 | QPP2 | QPP3 |
|---|---|---|---|---|---|---|---|
| Cx | 24 | 25 | 24 | | 23 | 25 | 24 |
| C | 25 | 26 | 26 | | 25 | ~~22~~ | 25 |
| T | 20 | 21 | 25 | | 24 | 27 | 22 |
| H | 24 | 19 | 24 | | 17 | 26 | 22 |
| A | - | - | 22 | | 18 | 23 | - |
| BD | 22 | 22 | 26 | | 24 | 27 | 21 |
| S | 24 | 24 | 22 | | 21 | ~~27~~ | 24 |

dash: number of voxels with significant peak less than 15

strikethrough: qualitatively no obvious second mode in the histogram

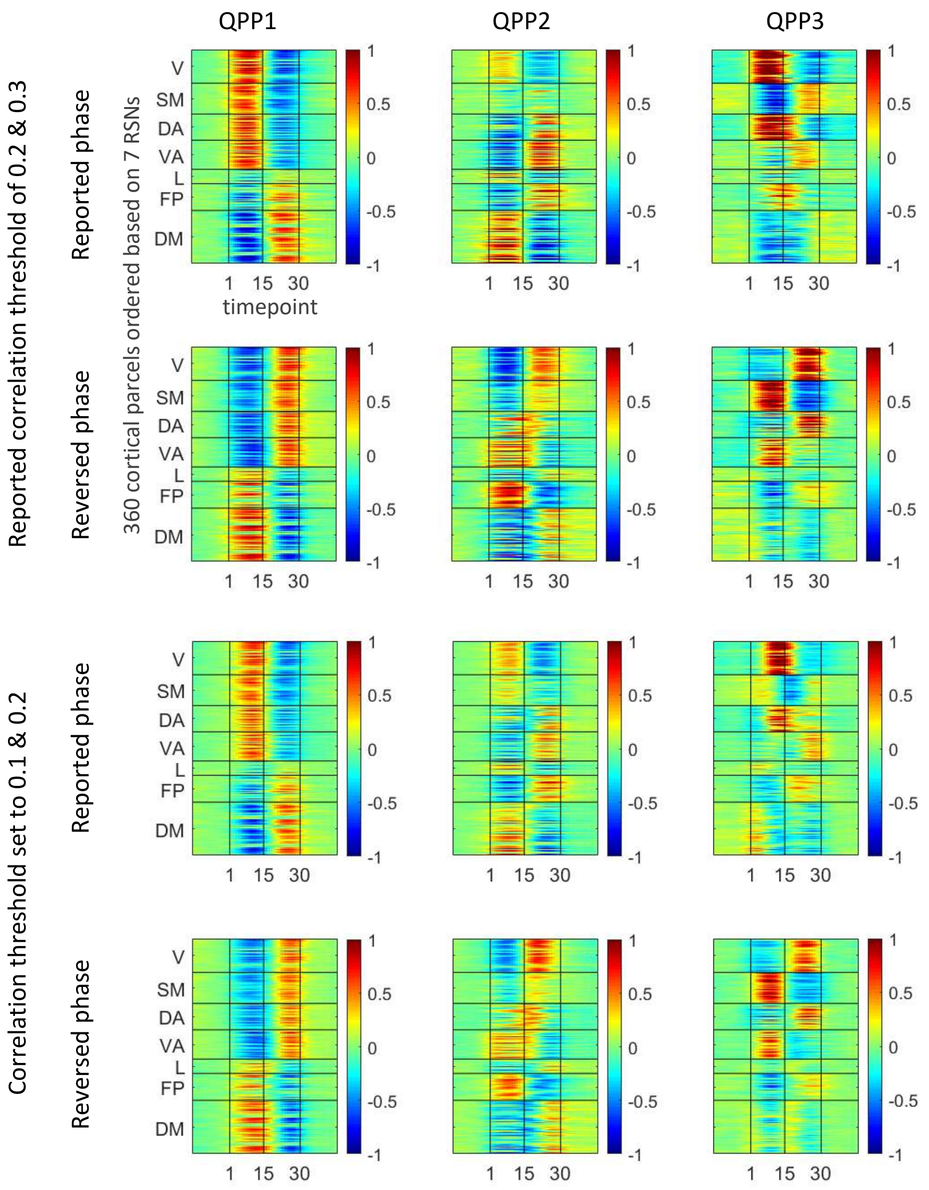


| Correlation threshold | Basic Metrics | QPP1 | QPP2 | QPP3 |
|---|---|---|---|---|
| 0.2 & 0.3 | Strength | 0.40±0.02 | 0.37±0.02 | 0.35±0.02 |
| | Occurrence intervals (s) | 51.3±6.2 | 64.6±9.4 | 78.5±14.1 |
| 0.1 & 0.2 | Strength | 0.35±0.03 | 0.30±0.02 | 0.27±0.01 |
| | Occurrence intervals (s) | 40.3±2.6 | 42.5±3.4 | 46.4±4.2 |

Fig.S26 Group QPPs (in parcel space) detected with two different correlation thresholds. First two rows are QPPs detected based on the setting of this report, which is the correlation threshold of 0.2 for the first two iterations of the QPP algorithm and 0.3 for the remaining iterations (shown as 0.2 & 0.3). Last two rows are QPPs detected based on the setting of all the prior work (0.1 for the first two iterations of the QPP algorithm and 0.2 for the rest of the iterations, shows as 0.1 & 0.2). In each subplot a group QPP in parcel-space is shown as a 2D array of 360 cortical parcels (reordered based on seven RSNs) by 30 timepoints (15 timepoints at each end for temporal extension, see Fig.S5). QPPs with the reversed phase are included here as another support for the previous supplementary analysis that showed robustness of QPP features with regards to the phase-related setting (Fig.S25). As qualitatively suggested by the figure, slight change of correlation threshold does not influence the detected QPPs. Quantitatively, the similarity between two patterns detected by the abovementioned correlation thresholds is 0.992, 0.926 and 0.843, the reported phase and 0.996, 0.960 and 0.933, the reversed phase, for QPP1, QPP2 and QPP3, respectively (similarity is quantified via the fine-phase matching described in Fig.S5). Lowering the correlation threshold in detecting a QPP, however, decreases the correlation of that QPP with its contributing segments (strength) but increases the number of its contributing segments; the latter is equivalent to a decrease in the occurrence interval between the contributing segments of that QPP. Basic metrics of QPPs brought in the table above indicate such changes.
